# Gene tree discord, simplex plots, and statistical tests under the coalescent

**DOI:** 10.1101/2020.02.13.948083

**Authors:** Elizabeth S. Allman, Jonathan D. Mitchell, John A. Rhodes

## Abstract

A simple graphical device, the simplex plot of quartet concordance factors, is introduced to aid in the exploration of a collection of gene trees on a common set of taxa. A single plot summarizes all gene tree discord, and allows for visual comparison to the expected discord from the multispecies coalescent model (MSC) of incomplete lineage sorting on a species tree. A formal statistical procedure is described that can quantify the deviation from expectation for each subset of four taxa, suggesting when the data is not in accord with the MSC, and thus either gene tree inference error is substantial or a more complex model such as that on a network may be required. If the collection of gene trees appears to be in accord with the MSC, the plots may reveal when substantial incomplete lineage sorting is present and coalescent based species tree inference is preferred over concatenation approaches. Applications to both simulated and empirical multilocus data sets illustrate the insights provided.

---

When analyzing multilocus data sets for a collection of taxa, it is common to begin by inferring *gene trees* for each locus. Assuming no recombination occurred within loci, trees should adequately describe their individual histories, regardless of the more complex relationships that may occur at the population level. When a large majority of the gene trees show the same relationships between the sampled loci for all taxa, it is generally accepted that those relationships indicate evolutionary relationships of the taxa, as depicted by a *species tree* or, more accurately, a *population tree*. Although the few gene trees that differ might be attributed to gene tree inference error, or to a small amount of incomplete lineage sorting (ILS), or to a combination of the two, an overwhelmingly dominant signal of a single tree tends to make the precise cause of the variation of less concern.

In practice, however, many multilocus data sets show substantial discord among gene trees. If the discord results primarily from gene tree inference error, due to sampling error from short loci, then a partitioned analysis of the concatenated sequences might be justified. If incomplete lineage sorting is the dominant source of discord, then a species tree method based on the *multispecies coalescent* (MSC) *model* should be preferred. If instead hybridization or lateral gene flow has led to discordant gene trees, then attempting to infer a species tree may be mistaken, as a network may provide better fit. A final possibility is that the sequence evolution models used to infer the gene trees fit poorly, and we have substantial gene tree inference error without any understanding of its form, and an improved gene tree analysis is essential to proceed.

Assessing the nature and cause of gene tree discord is a major challenge, for which many techniques are needed. A fundamental issue is the difficulty of directly grasping the similarities and differences across a large collection of gene trees. In this article we introduce a simple visualization approach that, in a single plot, can illustrate much about a gene tree collection, enough to suggest what further analysis might be appropriate. We describe several statistical tests associated to these plots, that, under the MSC model, can indicate poor fit of gene trees to a species tree. This might be due to either substantial gene tree inference error or to more complicated biological processes than those captured by the MSC on a species tree.

The plots introduced here are based on the counts of quartets (4-taxon subtrees) displayed on gene trees. Of course the use of quartets for describing trees and quantifying gene tree variation is far from novel. For instance, the relationship between quartets displayed on a tree and the full topological tree is an important topic in the combinatorial aspects of phylogenetics [Semple and Steel, 2005]. More recently, quartets on a collection of gene trees have played a role in methods for inferring species relationships, for both species trees [Larget et al., 2010, Zhang et al., 2018, Rhodes, 2019] and species networks [Solís-Lemus and Ané, 2016, Allman et al., 2019b]. Frequencies of quartets displayed on gene trees, called *concordance factors*, were introduced by Larget et al. [2010] and their expected form under the MSC model of ILS on a tree investigated by Allman et al. [2011]. Stenz et al. [2015] developed one approach to using quartet concordance factors for judging fit of gene trees to a species tree. However, the only previous visualization tool based on quartet counts that we are aware of was given by Sayyari et al. [2018], and simply uses a separate bar chart for each choice of four taxa to illustrate three frequencies.

The first novelty in our use of quartets is a single visualization that combines all quartet concordance factors for all choices of four taxa. This gives a comprehensive view of the discordance across the entire gene tree data in a single picture, called a *simplex plot*. Even with no further formal analysis this plot can provide substantial insight into a data set, once a researcher has learned to interpret it. We also describe statistical tests that can quantify the strength with which one might draw conclusions from the plots. We give a thorough development of quartet concordance factor simplex plots, as we believe these should become a standard means of summarizing and communicating information about multilocus data sets. Implementations in R of all methods presented here are available in the MSCquartets package [Allman et al., 2019a], available for download from CRAN.

## Methods

In this section, we first recall the notion of a *quartet concordance factor* [Larget et al., 2010] and describe the simplex plot for its visualization. After discussing expected concordance factors under several models, we describe statistical tests for the MSC model based on them. This is followed by illustrations of plots and tests using simulated data of both gene trees sampled from the MSC, and inferred gene trees from sequences simulated on the sampled gene trees.

To simplify language, we initially refer to a collection of gene trees as *data*. If the trees arose from a simulation process of the MSC, this language is entirely appropriate. If, however, they were inferred from empirical sequence data, this is a misnomer, as the gene trees are not raw observations, but rather obtained from processing the observed sequences. When we turn to considering simulated gene trees, and gene trees inferred from simulated sequences, we more carefully indicate that gene trees are not raw data.

### Quartet Concordance Factors and Simplex Plots

Consider a data set of unrooted topological gene trees relating *N* taxa. If gene trees are rooted or metric, we simply ignore that additional information. Trees in the data set may have ‘missing taxa’ and thus have fewer than *N* leaves. For any choice of four of the taxa *a, b, c, d*, each of the gene trees on which all four appear displays some unrooted *quartet tree* relating them. The possibilities for this quartet tree are the three resolved topologies, denoted *ab|cd*, *ac|bd*, and *ad|bc*, and the unresolved, or star, topology, denoted *abcd*. Tabulating the number of occurrences of each across the gene trees, we obtain four counts, 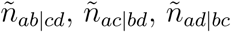, and 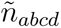.

Since star quartets on gene trees are generally viewed as representing soft polytomies (ignorance of a true resolution) rather than hard ones (historical truth), and furthermore have probability 0 under the MSC, we must choose how to treat them. With no extra information, there are two defensible options. One is to simply discard the count 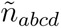 of star quartets, reducing the data for these four taxa to only those trees displaying a resolved quartet. Another is to consider a star quartet as contributing a count of 1/3 to each of the resolved topologies, adding 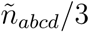 to each of the other counts. As lack of resolution in gene trees may arise in several ways, the choice between these should be based on an understanding of the gene data. Polytomies due to suspected short edges in the true gene tree (i.e., true ‘near polytomies’) might be better handled by assigning 1/3 to each of the resolved topologies, while those reflecting total lack of information are better discarded. We emphasize that by describing an edge as ‘short’ on a gene tree we mean it is short when measured in substitution units, which are not the standard units on species trees. A short species tree edge, in coalescent units, may be due to either a duration of a small number of generations, or a large population size, or a combination of these factors, yet a sufficiently large substitution rate may still lead to long edges in gene trees.

The three resultant counts of resolved quartets constitute the *quartet count concordance factor* (qcCF) for taxa *a, b, c, d*, denoted by

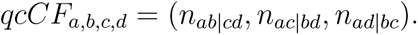

If some of the taxa are missing from some of the gene trees, or some unresolved quartets have been discarded, then the total count

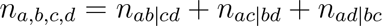

may vary with the choice of the four taxa, and be less than the number *n* of gene trees in the data set.

Normalizing *qcCF* so the sum of entries is 1 gives the *empirical quartet concordance factor* for *a, b, c, d*,

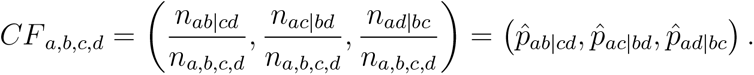

The three entries are nonnegative, and should be interpreted as estimates of the probabilities that a randomly chosen gene tree displays the resolved quartets on *a, b, c, d*.

Geometrically, *CF*_*a,b,c,d*_ is a point in 3-dimensional space, which lies on the plane *x* + *y* + *z* = 1 and in the octant where *x* ⩾ 0, *y* ⩾ 0, *z* ⩾ 0. The set of all such points (*x, y, z*) forms a 2-dimensional equilateral triangle, as shown in Figure 1a, called the 2-dimensional *probability simplex*. This simplex is more conveniently drawn as its isometric image in the plane, as in Figure 1b. (Formulas for the mapping are given by Mitchell et al. [2019].) The vertices are still labelled (1, 0, 0), (0, 1, 0), and (0, 0, 1) to emphasize the source of the triangle.

**Fig. 1.**
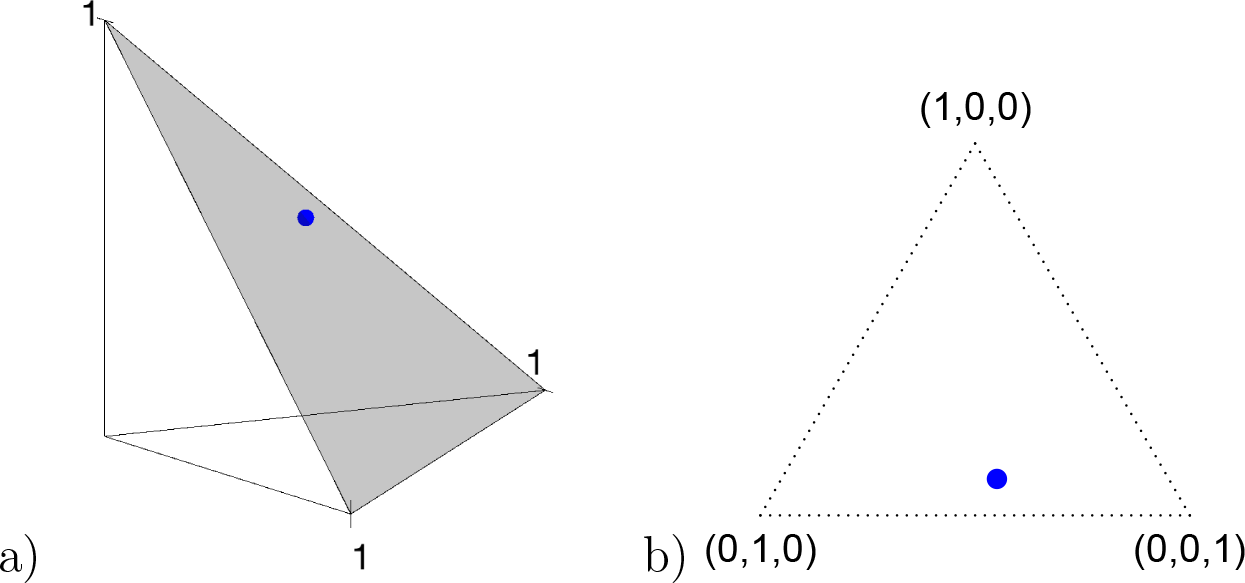
a) The probability simplex in 3-dimensional space is a 2-dimensional equilateral triangle. Any quartet concordance factor can be viewed as a point in the simplex. b) The simplex, and a concordance factor depicted in it, can be drawn in the plane.

Any quartet concordance factor is represented by a point in this triangle. The vertices represent *CF*s where across all gene trees only one quartet topology occurs for the four taxa, the edges of the simplex include those *CF*s where only two quartet topologies occur, and interior points those *CF*s where all three topologies occur. The centroid (1/3, 1/3, 1/3) is the *CF* with equal frequencies of the three topologies.

Note further that if the four taxa *a, b, c, d* are reordered, the entries of the *CF* are reordered as well, which gives up to 6 symmetrically placed points that might represent essentially the same information. For now, we merely insist on having chosen some specific order for each set of four taxa.

Simplex plots of this sort (also called ternary plots) are used in a variety of fields. For example, they can represent color palettes obtained by mixing three primary colors. In phylogenetics they have been used for different purposes than described here, by Strimmer and von Haeseler [1997] and Smith [2019]. Their first published use for representing quartet concordance factors was in several works closely related to this one [Baños, 2019, Mitchell et al., 2019, Allman et al., 2019b].

### Model Predictions for Concordance Factors

While simplex plots of quartet concordance factors can be made from any collection of gene trees, an interpretation of them hinges on modeling the process by which the trees were generated. Here we explore *expected quartet concordance factors* under two models of gene tree generation on a species tree.

Consider first a model for which all gene trees are topologically concordant with the species tree, a model we refer to as the *no discord* (ND) *model*. This might also be called the *concatenation model* as it posits that in the absence of any gene tree inference error there is complete gene tree concordance, the only model-based viewpoint which provides justification for concatenation approaches to species tree inference.

Under the ND model, regardless of the choice of four taxa, the expected quartet concordance factor is one of (1, 0, 0), (0, 1, 0), or (0, 0, 1), depending on the arbitrary ordering chosen for the taxa. Thus a simplex plot of all expected *CF*s under the ND model shows points plotted only in the vertices of the simplex, as in Figure 2a. While particular choices of orderings for the different 4-taxon subsets might lead to points plotted at only 1 or 2 of the vertices, random orderings typically lead to points at all 3. Regardless, no expected *CF*s will appear except at the vertices.

**Fig. 2.**
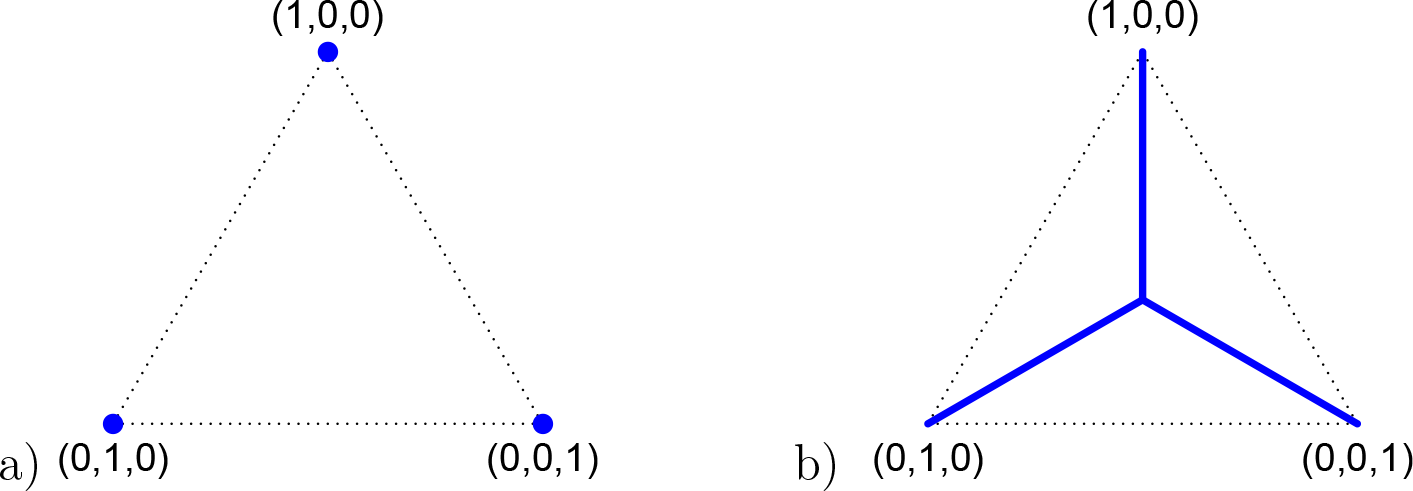
a) A simplex plot showing possible expected *CF*s under the ‘no discord’ ND model when no specific species tree is hypothesized. The expected *CF*s may appear only at the vertices of the simplex. b) A simplex plot showing possible expected *CF*s under the MSC model of ILS when no specific species tree is hypothesized. The *CF*s may appear at any point on the three line segments from the centroid (1/3, 1/3, 1/3) to the vertices, forming model T3. More precisely, the *CF*s will be a finite set of points on these lines, whose locations depend on the branch lengths on the species tree.

Now consider the MSC model on some metric species tree *σ*. As shown by Allman et al. [2011], the expected *CF* for a set of four taxa is, depending on taxon ordering, one of (*p, q, q*), (*q, p, q*), or (*q, q, p*), with

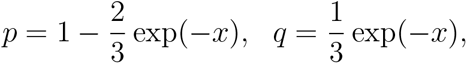

where *x* is the length in coalescent units of the internal edge on the species quartet tree for the chosen four taxa. The largest entry, *p*, in the expected *CF* indicates the correct topology for the induced quartet species tree. For example, if the tree *ab|cd* is displayed on the species tree, then the expected *CF* is (*p, q, q*). Since *p* ⩾ 1/3 ⩾ *q*, in a simplex plot the expected *CF* lies on one of the three interior line segments shown in Figure 2b. These line segments extend from the centroid of the simplex, (1/3, 1/3, 1/3), to the three vertices.

An expected *CF* at the centroid arises only when the quartet on the species tree is unresolved, leading to equal probabilities of each of the resolved gene tree quartets. Expected *CF*s on the vertical segment arise when the species tree quartet displays the topology *ab|cd*, with longer internal edge lengths on the species quartet tree moving the point toward the vertex (1, 0, 0). That vertex itself corresponds to an infinite edge length, which results in no expected discord between gene trees and the species tree. The other line segments similarly contain expected *CF*s for the other two species quartet topologies. Following Mitchell et al. [2019], we refer to the model in which the expected *CF* for every quartet lies on the line segments in Figure 2b as *model T3*, which stands for ‘any of <u>3</u> possible species quartet trees’. If gene tree production is modeled by the MSC model on *any n*-taxon species tree, regardless of topology or branch lengths, the expected *CF*s lie in the model T3.

If the unrooted topology *S* of the metric species tree *σ* is known, or more realistically hypothesized, the four taxa can be ordered so the first entry of their *CF* corresponds to the quartet displayed on *S*. Then under the ND model the expected *CF* for every choice of four taxa will be (1, 0, 0), lying at the top vertex of the simplex. Under the MSC model on *σ*, the expected *CF*s all lie on the vertical line from the centroid (1/3, 1/3, 1/3) to the top vertex. Again following Mitchell et al. [2019], we refer to this as *model T1*, which stands for ‘<u>1</u> specific species quartet <u>t</u>ree’. Figure 3 shows these possibilities graphically.

**Fig. 3.**
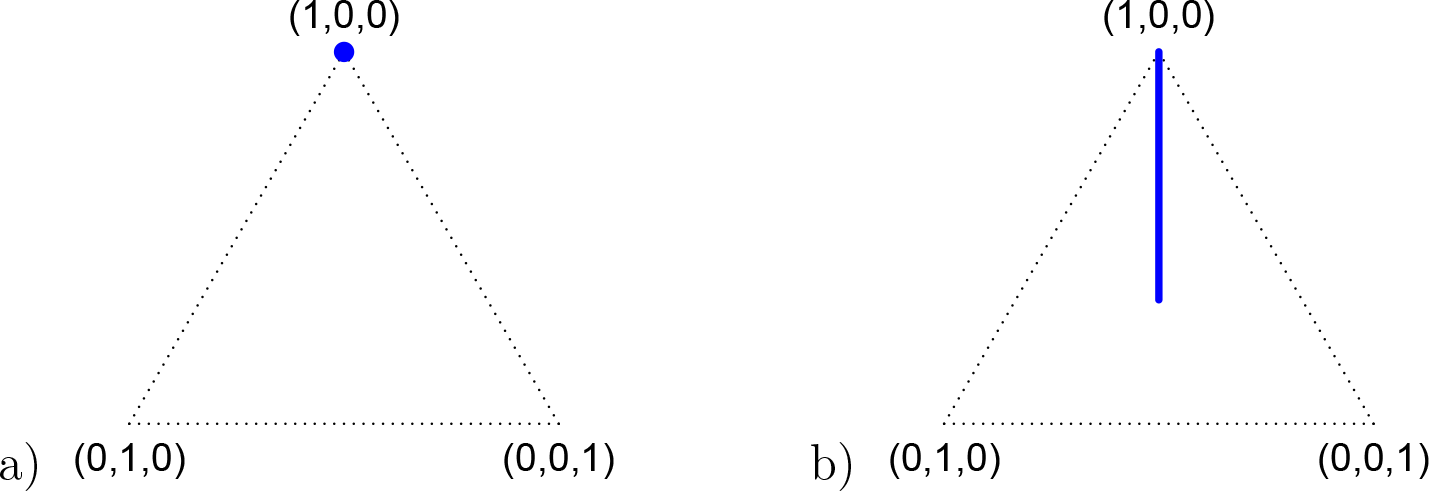
a) A simplex plot showing expected *CF*s under the ‘no discord’ ND model with the correct species tree topology hypothesized. The expected *CF*s may only appear at the top vertex of the simplex. b) A simplex plot showing all possible expected *CF*s under the MSC model of ILS when the correct species tree topology is hypothesized. The expected *CF*s may appear at any point on the vertical line segment from the centroid (1/3, 1/3, 1/3) to the top vertex, forming model T1. More precisely, the expected *CF*s will be a finite set of points on this line, whose locations depend on the branch lengths on the species tree.

For a fixed metric species tree *σ* on *N* taxa, each of the 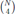 expected *CF*s can be plotted on the same simplex plot. Because under the MSC the expected *CF* depends only on the internal branch length of the quartet tree on *σ*, some of these points will be the same; the internal quartet branch is determined by its endpoints, which must be two of the *N* − 2 internal nodes on an unrooted binary tree. This means there can be only 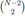 different expected *CF*s, up to ordering of the entries. Thus, up to the ordering, there are on average *N* (*N* − 1)/12 copies of each expected *CF*. For a simplex plot made under model T3, these are typically scattered across the model line segments, and some may appear symmetrically located. In a T1 simplex plot made by hypothesizing the correct species tree topology, the expected *CF*s will all lie on the same vertical line segment. Figure 4 illustrates this for the expected *CF*s arising under the MSC for a specific species tree.

**Fig. 4.**
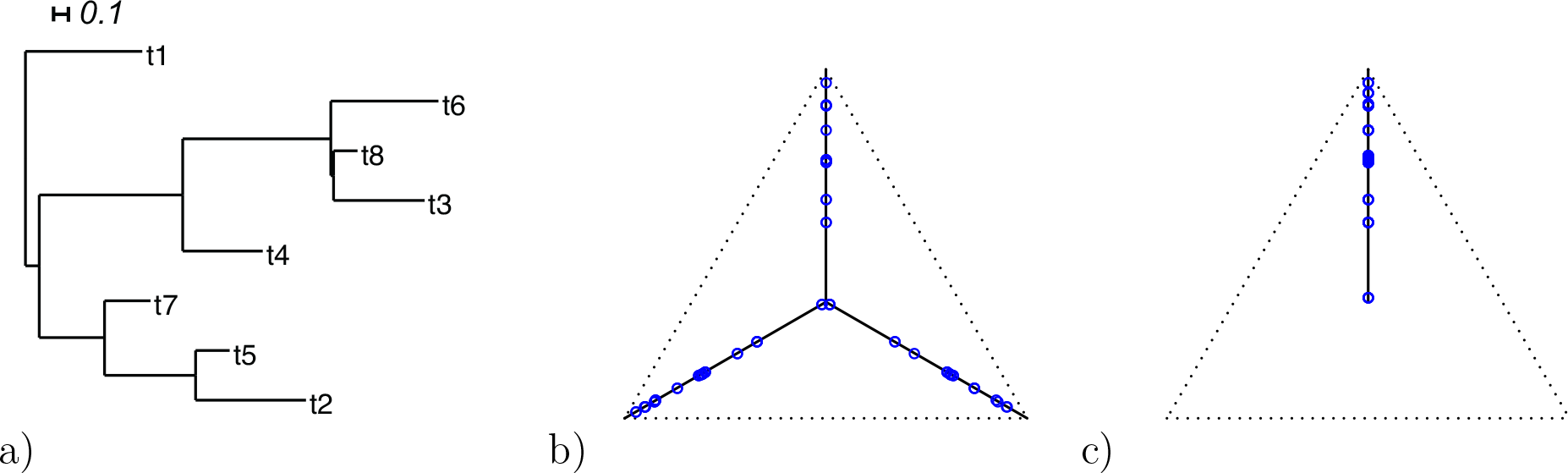
a) An 8-taxon species tree, with branch lengths in coalescent units. b) A simplex plot showing expected *CF*s under the MSC on this tree, for the model T3, in which the topology of the species tree is not known. The expected *CF*s appear at a discrete set of points along the three line segments, with some symmetrically located. c) A simplex plot showing expected *CF*s under the MSC when the correct species tree topology has been hypothesized, so the model T1 applies. The expected *CF*s appear at a discrete set of points on the vertical line segment from the centroid (1/3, 1/3, 1/3) to the top vertex. As these two plots were constructed from expected *CF*s for the same species tree, figure (c) can be obtained from (b) by ‘reflecting’ all points on the three lines onto the vertical one. Which line segment a point in (b) appears on is determined by an arbitrary ordering of the entries of the *CF*. The species tree with branch lengths in coalescent units is (t1:0.75,(((t6:0.69,(t8:0.15,t3:0.58):0.02):0.95,t4:0.51):0.92,(t7:0.29,(t5:0.22, t2:0.71):0.58):0.42):0.09).

It is instructive to modify the tree in Figure 4a to see the effect on expected *CF*s. In Figure 5, we show simplex plots of expected *CF*s for such a modified tree, which differs only by the introduction of a population bottleneck on a single edge. As lengths in coalescent units are the number of generations divided by population size, this is modeled by simply increasing that edge length. Many expected *CF*s are left unchanged, because at least three of the four taxa lie on one side of this edge. However, if two of the four taxa lie on each side of the edge, there is an impact on the *CF*. For a tight bottleneck, essentially all the gene trees will display quartet trees for these taxa that match those of the species tree. The expected *CF*s under the MSC then lie close to the vertices of the simplex. Indeed, in Figure 5b,c we see fewer expected *CF*s far from the vertices than in Figure 4b,c. Note that under the ND model we would see no change in the simplex plot when a bottleneck was introduced.

**Fig. 5.**
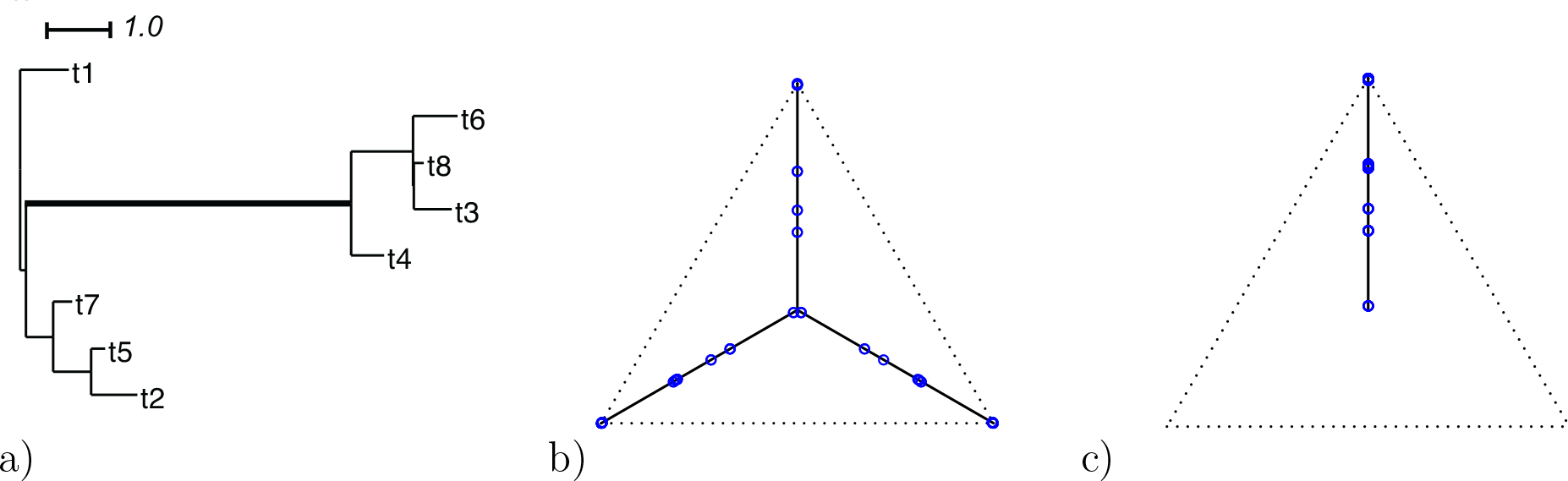
a) A species tree identical to that in Figure 4a except for a population bottleneck causing one branch to be much longer. Simplex plots showing expected *CF*s under the MSC on this tree under b) the model T3, and c) the model T1 with the correct species tree topology hypothesized. The bottleneck shifts certain expected *CF*s to appear at the simplex vertices. The species tree with branch lengths in coalescent units is (t1:0.75,(((t6:0.69,(t8:0.15,t3:0.58):0.02):0.95,t4:0.51):**5**,(t7:0.29,(t5:0.22, t2:0.71):0.58):0.42):0.09). The changed branch length is shown in bold.

For any model of gene tree generation more general than the MSC, such as one including hybridization, or gene duplication and loss, etc., expected *CF*s may not lie on the T3 or T1 model lines. An example of this, given by introducing a single hybridization into the tree of Figure 4, is shown in Figure 6b,c for the Network MSC, where different colors and shapes indicate whether expected *CF*s lie on or off the model lines. For this example the species tree used for Figure 4 is modified to the network of Figure 6a with one hybrid node which has only one descendant taxon. Because of this simple structure, one can also view the gene tree distribution under this network model as a mixture of those that arise on two species trees under the MSC, where the trees differ by a single subtree-prune-and-regraft move of a pendant edge. While only *CF*s involving the single hybrid taxon are moved from their locations in the original plot, many *CF*s are affected, often displaced from the model lines. In general, the number affected depends on the number of edges in the cycle introduced in the network, the number of descendants of the hybrid node, and other details of the network structure. In contrast, the situation is much simpler for the ND model on this network, as is shown in Figure 6d,e, where *CF*s lie either at vertices, or at points on a edge 30% of the way between vertices.

**Fig. 6.**
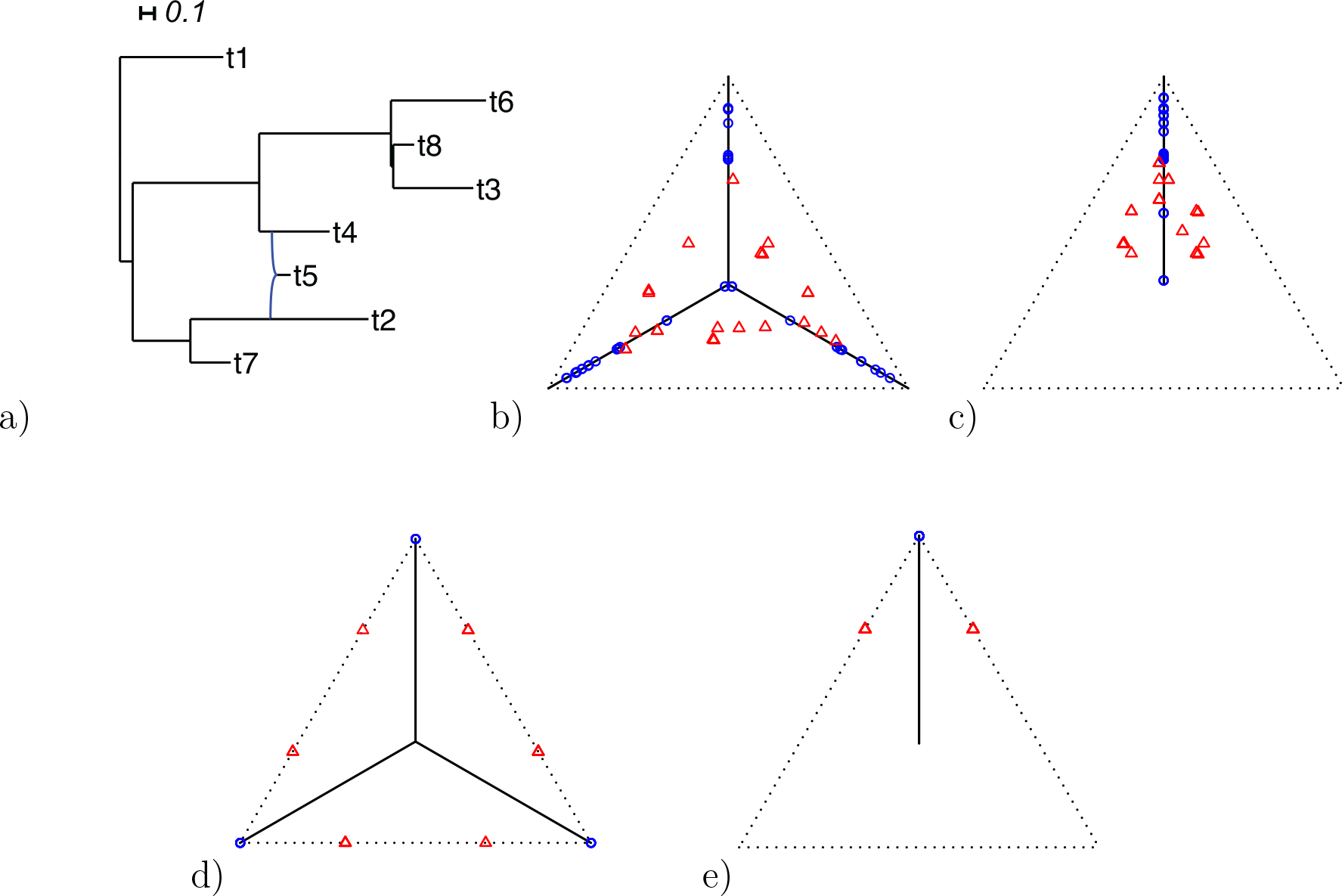
a) A hybridization network obtained from the tree of Figure 4a by the introduction of an additional edge. Simplex plots show expected *CF*s under the Network MSC for b) the model T3, and c) the model T1 with the species tree of Figure 4a hypothesized. This shifts certain expected *CF*s off the T3 or T1 tree model lines. Simplex plots under the Network ND model are shown in d) and e). Red triangles mark expected *CF*s off the model lines, while blue circles mark those on the lines. The gene tree distribution here is a mixture with 70% of gene trees arising from the MSC on the species tree in Figure 4a, and 30% from the MSC on the species tree (t1:0.75,(((t6:0.69,(t8:0.15,t3:0.58):0.02):0.95,(t4:0.42,t5:0.21):0.09):0.92,(t7:0.29,t2:1.29):0.42):0.09), both of which are displayed in the network.

Allman et al. [2019b] showed that even if hybridization is limited to that described by networks that are level-1, then an expected *CF* under the Network MSC may be located at any point in the simplex. Thus while these simplex plots may visually suggest whether the ND or the MSC models might give reasonable fit to a data set, they are not able to immediately suggest which of many more complex biological processes must be modeled to capture important features of some data sets.

### Sampling Error and Statistical Tests

Now we consider empirical concordance factors from a finite sample of gene trees drawn from a model. If a model allows for gene trees to vary, a finite sample is unlikely to match expectations due to *sampling error*. However, statistical tests can be developed to measure whether the deviation from expectation is sufficiently large for a given data set to reject the model as a likely source of the data. Note that this sampling error is distinct from the *gene tree inference error*, which arises when the gene trees are inferred from sequence data.

*The ND model* Under the ND model, every gene tree in a sample matches the species tree topology. Thus with probability 1 empirical *CF*s for all choices of four taxa must lie at the vertices of the simplex. This is as shown in Figure 2a for the model T3, or as in Figure 3a for the model T1 if the correct species tree topology is hypothesized. There is no possibility of sampling error causing empirical *CF*s to vary from expectation.

A procedure using a *qcCF* as a statistic for testing the ND model is then trivial. For a fixed choice of four taxa, null and alternative hypotheses are:

*H*_0_ : The gene trees arose from the ND model on a species tree.
*H*_1_ : The gene trees did not arise from the ND model on a species tree.

*H*_0_ is then rejected exactly when an empirical *CF* lies anywhere other than at a vertex in a T3 simplex plot.

For a specific hypothesized unrooted species tree topology *S*, the null and alternative hypotheses are:

*H*_0_ : The gene trees arose from the ND model on a species tree with topology *S*.
*H*_1_ : The gene trees did not arise from the ND model on a species tree with topology *S*.

Then *H*_0_ is rejected if the T1 simplex for *S* shows an empirical *CF* at any point other than the top vertex. If these tests are applied for all choices of four taxa, the ND model on an unknown tree, or more specifically on trees with topology *S*, is rejected if it is rejected for any single choice of four taxa.

Of course such a test has only pedagogical value. In practice, gene trees should not be viewed as sampled from the ND model, but rather from a model with inference error introduced when gene trees are inferred from sequence data. Such an error model would ideally account for impact of sampling error in the sequences (since they are of finite length) as well as any mild model misspecification in the substitution processes used in inference. Use of the ND model hinges on a belief that the observed discord all arises from such error. Unfortunately, no comprehensive error model has been proposed, and little work has been done to even begin to quantify such error. Without such a model, no more useful test can be developed.

The *MSC model* For the MSC model, in contrast, it is possible to develop useful statistical tests to apply to *qcCF* statistics. Focusing first on the T1 simplex plot, assuming the correct species tree topology *S* is hypothesized, the expected *CF* is a point on the vertical line segment shown in Figure 3b. Then a finite sample of *n*_*a,b,c,d*_ gene quartets has quartet counts determined by a random sample from the trinomial distribution with parameters given by the expected *CF*. (If an unresolved quartet count has been treated as the non-integer counts (1/3,1/3,1/3), then technically the *qcCF* cannot be viewed as a trinomial sample, but this raises no practical issues.) An empirical *CF*_*a,b,c,d*_, computed from a finite sample of gene trees, should then be a point near the expected *CF*, and as the sample size *n*_*a,b,c,d*_ is increased, by the Law of Large Numbers it is increasingly probable that it lies close to the expected *CF*. Informally, if an empirical *CF*_*a,b,c,d*_ is near the vertical line segment it offers support for *S*, while if it is far away it suggests rejection of *S*. However, the size of the sample must be taken into account in giving precise meanings to ‘near’ and ‘far’.

To formalize these ideas, for four taxa and an unrooted species tree topology *S* consider the null and alternative hypotheses:

*H*_0_ : The gene trees arose from the MSC model on a species tree with topology *S*.
*H*_1_ : The gene trees did not arise from the MSC model on a species tree with topology *S*.

For four chosen taxa, compute the *qcCF* from the gene tree data. Then from the *qcCF*, compute maxima of the likelihood functions under both the null and alternative models in order to find the likelihood ratio statistic for the *qcCF*.

A standard approach, suggested by Degnan and Rosenberg [2009], would be to use a *χ*^2^ distribution with 1 degree of freedom to obtain a *p*-value. While this can be justified if the *CF* is not near the centroid or a vertex of the simplex, it is problematic in some regions. The theoretical argument for the use of the *χ*^2^ distribution requires the model to be well approximated by its tangent line, and near the boundary point of the T1 model at (1/3, 1/3, 1/3) this is not valid. Near the vertex (1, 0, 0), which corresponds to a long internal branch on the species tree quartet, we may have very small expected counts of the discordant quartet topologies, and other well-known issues arise with using the *χ*^2^ when expected counts are low.

When not hypothesizing a specific species tree topology, a test is performed for the model T3. Again, far from the vertices and the centroid a standard test can be formulated with the likelihood ratio statistic using the *χ*^2^ distribution with 1 degree of freedom. However, the singularity of the model, where the three lines come together, prevents justification of this in its vicinity. The possibility of small expected counts near the vertices also remains problematic.

Better behaved statistical tests for fit of *qcCF*s to the T1 and T3 models, which take into account the boundary or singularity at the centroid, have been developed by Mitchell et al. [2019], and are implemented in the R package `MSCquartets` [Allman et al., 2019a]. The package enables tabulation of all *qcCF*s for a set of gene trees, computation of *p*-values measuring fit to either of the models T1 or T3, and creation of a simplex plot. A small *p*-value indicates it is unlikely a *qcCF* arose under the model, suggesting rejection of the model. There are several options as to the precise test one might use, as discussed more fully in the documentation to the `MSCquartets` package and the paper [Mitchell et al., 2019]. In this work we adopt the default choices of the function `quartetTreeTest` in the package, which involves comparison of the statistic to a theoretical distribution in most cases, and an approximation derived from a precomputed bootstrap when expected discordant counts are small. This approximation is very accurate when total counts exceed 30, and allows for fast computations, which is essential when the tests are performed on the large number of *qcCF*s which arise when the number of taxa is large.

Care needs to be taken when using model T1 if the species tree topology is not chosen a priori. If an inferred species tree is chosen, gene trees may fit better than they would for the true species tree. In particular, the hypothesis test here is not designed to choose among several trees for which best fits the data.

Although the gene trees may have many taxa, the tests described so far are performed for a single *qcCF*, for one choice of four taxa, at a time. When performed for each choice of four taxa, this raises issues with multiple comparisons being performed on the same data set (the sample of gene trees). Moreover, the observed gene tree quartets are not independent of one another, and their precise dependence structure has not yet been fully analyzed.

To address the multiple testing problem, one can apply the Holm-Bonferroni method [Holm, 1979], which does not require independence of the hypotheses for the tests. Since performing a large number of tests makes it more likely some statistics will be extreme even under the null hypothesis, the method can be viewed as a form of inflation of the *p*-values, to reduce the probability of a Type I error (erroneous rejection of the null hypothesis). For a significance level *α*, the Holm-Bonferroni method ensures the probability of one or more Type 1 errors is at most *α*. The method provides a conservative means of rejecting only a subset of the hypotheses for individual quartets that might otherwise have been rejected. When applied to all *qcCF*s for a collection of gene trees, it thus enables one to pinpoint quartets whose tree-likeness is doubtful for the T3 model, or which significantly deviate from the hypothesized tree for the T1 model. This method is also implemented in `MSCquartets`.

An important point for interpreting simplex plots and hypothesis test results is the difference between a *qcCF*, the counts of quartet topologies, and its associated empirical *CF*, the relative frequencies. Although the test is performed on a *qcCF* and thus includes information on the sample size, a simplex plot shows the empirical *CF*, which contains no information on sample size. Thus two *qcCF*s might lead to the same empirical *CF* plotted in a simplex, yet produce very different *p*-values under the same test, due to their differing sample sizes. If the sample size is small one might obtain a relatively large *p*-value, while a much larger sample size would give a small *p*-value.

In particular, when considering simplex plots of empirical *CF*s for different data sets, what is considered near or far from the model lines may change. If all taxa are present on all gene trees in both data sets, but the number of gene trees in the data sets differ, the *p*-value for a *qcCF* that gives the same point in both simplex plots with be larger for the smaller data set.

Even within a simplex plot for a single data set, if some taxa are missing from some gene trees then different choices of four taxa may lead to differing sample sizes. Thus a plotted empirical *CF* which had a large sample size may be associated to a small *p*-value, while another empirical *CF* plotted further from the model lines may produce a large *p*-value due to a smaller sample size. For a data set with some taxa missing from many gene trees, while other taxa are on most gene trees, this may complicate interpretation of a single simplex plot. However, if there are few missing taxa, or a uniform pattern of missing taxa, the effect will be negligible.

### Simulations

To illustrate these ideas, and the impact of gene tree inference error, we give a series of results from simulations, performed using `SimPhy` [Mallo et al., 2016] for the multispecies coalescent process and `Seq-Gen` [Rambaut and Grass, 1997] for sequence simulation, with gene trees inferred by IQ-TREE [Nguyen et al., 2015]. We briefly describe the simulation process here; full details can be found in the online appendix.

For all simulations we present, we fix the 15-taxon ultrametric species tree shown in Figure 7, with branch lengths in generations. With 15 taxa, there are 1365 *qcCF*s summarizing any gene tree collection.

**Fig. 7.**
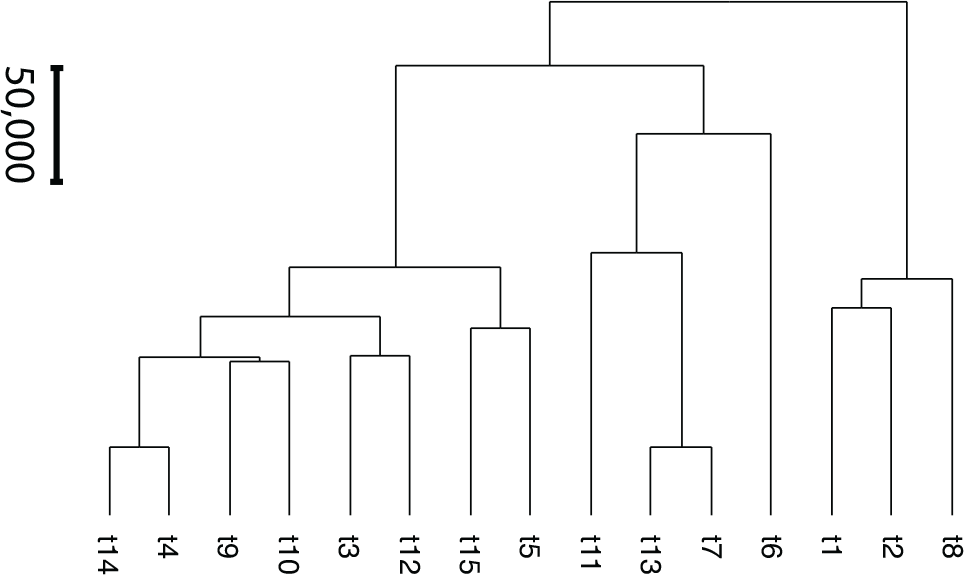
The species tree, with edge lengths in generations, used for simulations. See online Appendix for Newick description.

Since under the MSC the prevalence of ILS increases with population size *N*, we consider four sizes, *N* = 2, 5000, 10000, and 20000, which are held constant across all edges of the tree. The choice of *N* = 2 (the minimum allowed by `SimPhy`) leads to essentially instantaneous coalescence of lineages in a population and thus, up to minuscule error in branch lengths, gives gene trees from the ND model. Adopting a choice of parameters for a GTR model of sequence evolution from an empirical study, we considered conversion factors of *ν* = 10^*a*^ for *a* = −7, −6, −5 to convert gene tree edge lengths from number of generations to substitution units, a range progressing from little mutation to high levels. Of these, the central value *a* = −6 gives levels of substitution that are moderate, with the more extreme choices useful for bracketing behavior.

For each choice of *N* and *ν* we simulated 1000 gene trees under the MSC, giving samples of fully resolved gene trees without inference error. We then simulated sequences of length 500 bp along each gene tree. We next inferred maximum likelihood gene trees from these sequences, using a setting that caused all inferred edges of length ⩽ 10^−6^ to be contracted to length 0. This gave us samples of gene trees with inference error, which included some polytomies.

We investigate these simulated data sets in several ways. First, in Figure 8, for each value of *N* we plot expected *CF*s and empirical *CF*s from the gene tree sample without inference error. As the value of *ν* is irrelevant for this (other than the fact that our pipeline produced an independent sample of topological trees for each *ν*), we show such plots for only one 1000 tree sample, along with subsamples of 100 and 500 trees.

**Fig. 8.**
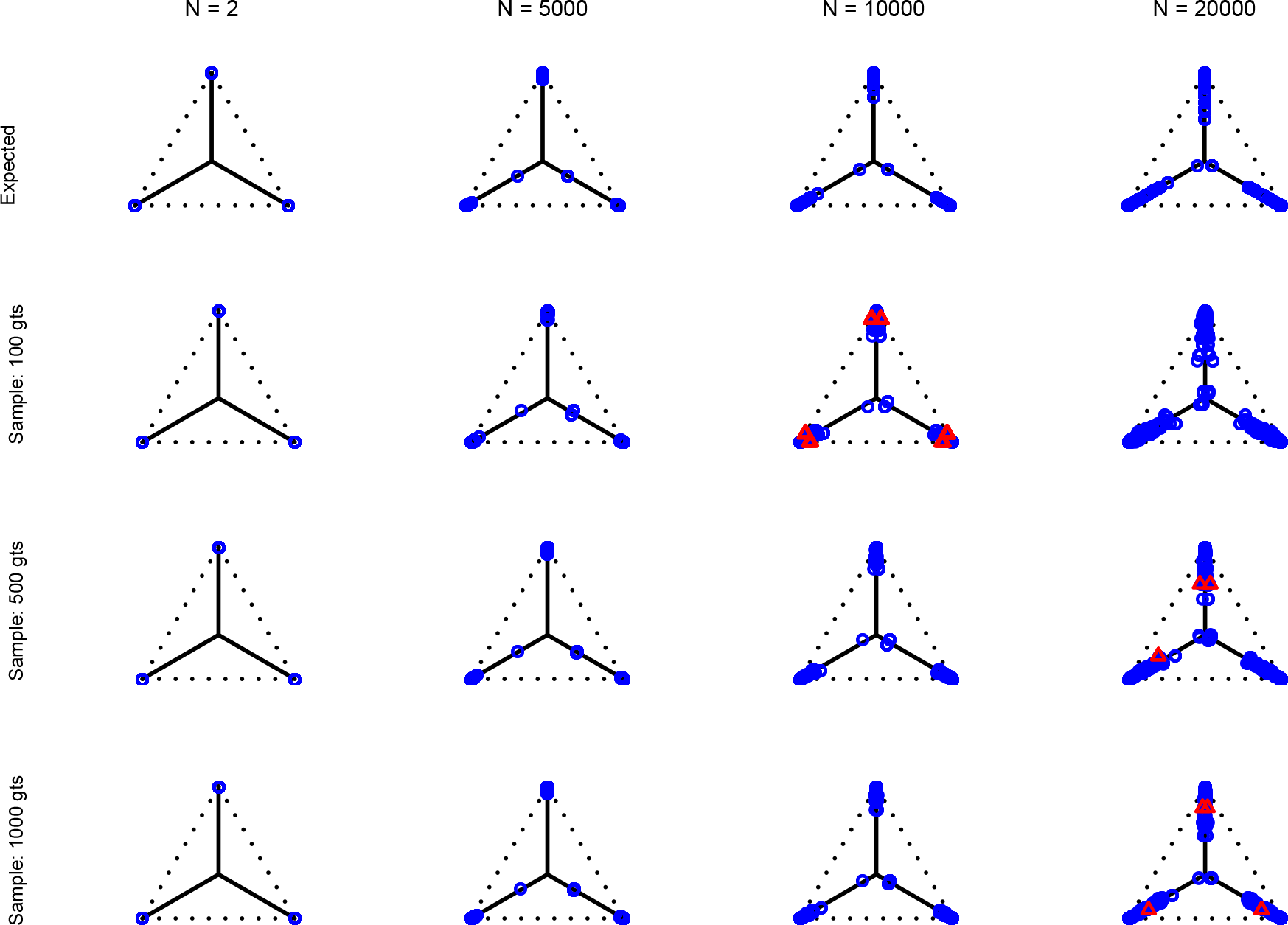
Row 1 shows expected *CF*s for the MSC on the species tree of Figure 7 for population sizes *N* = 2, 5000, 10000, and 20000. Rows 2,3,4 shows empirical *CF*s for samples of 100, 500, and 1000 gene trees with the same population sizes, simulated under the MSC by `SimPhy`. Red triangles denote empirical *CF*s for which the hypothesis test with level *α* = 0.01 rejects the T3 model, and thus the MSC on a tree.

In the top row of Figure 8 the expected *CF*s under the MSC for the various population sizes are shown. For population size *N* = 2 the plot appears to be identical to Figure 2a as the expected level of ILS is graphically imperceptible. Larger values of *N* produce more ILS, hence more discordant gene trees, and thus expected *CF*s move closer to the centroid. The bottom three rows of Figure 8 show empirical *CF*s from MSC samples of 100, 500, and 1000 gene trees. While close to their expectations, many empirical *CF*s lie off the model lines. However, as the number of gene trees is increased, the spread from expectations is reduced. Only MSC sampling error is present in the plots in the bottom rows, since the gene trees were simulated under the MSC process, and not inferred from sequences. Although we applied the T3 hypothesis test with level *α* = 0.01 to each *qcCF*, there were few rejections, as expected.

Next, Figure 9 shows simplex plots for the 1000 gene trees inferred from simulated sequences, for each value of *N* and *ν*, where counts of unresolved quartets are omitted. Similar figures for 100 and 500 gene trees with inference error, and for the same trees when unresolved quartets are redistributed among the resolved, are shown in the online appendix. While similar overall to both the expectation plots and the MSC sample plots of Figure 8, there are several differences to be noted. First, the cloud of *CF*s spreads further out from the model lines, and there are more rejections under the T3 test. Second, we see *CF*s moved inward slightly from the vertices, toward the centroid. Both of these changes are reasonable since the use of inferred gene trees has introduced additional error. Assuming that error has no topological bias it could plausibly produce a greater spread and a reduction in the most frequent quartet topology.

**Fig. 9.**
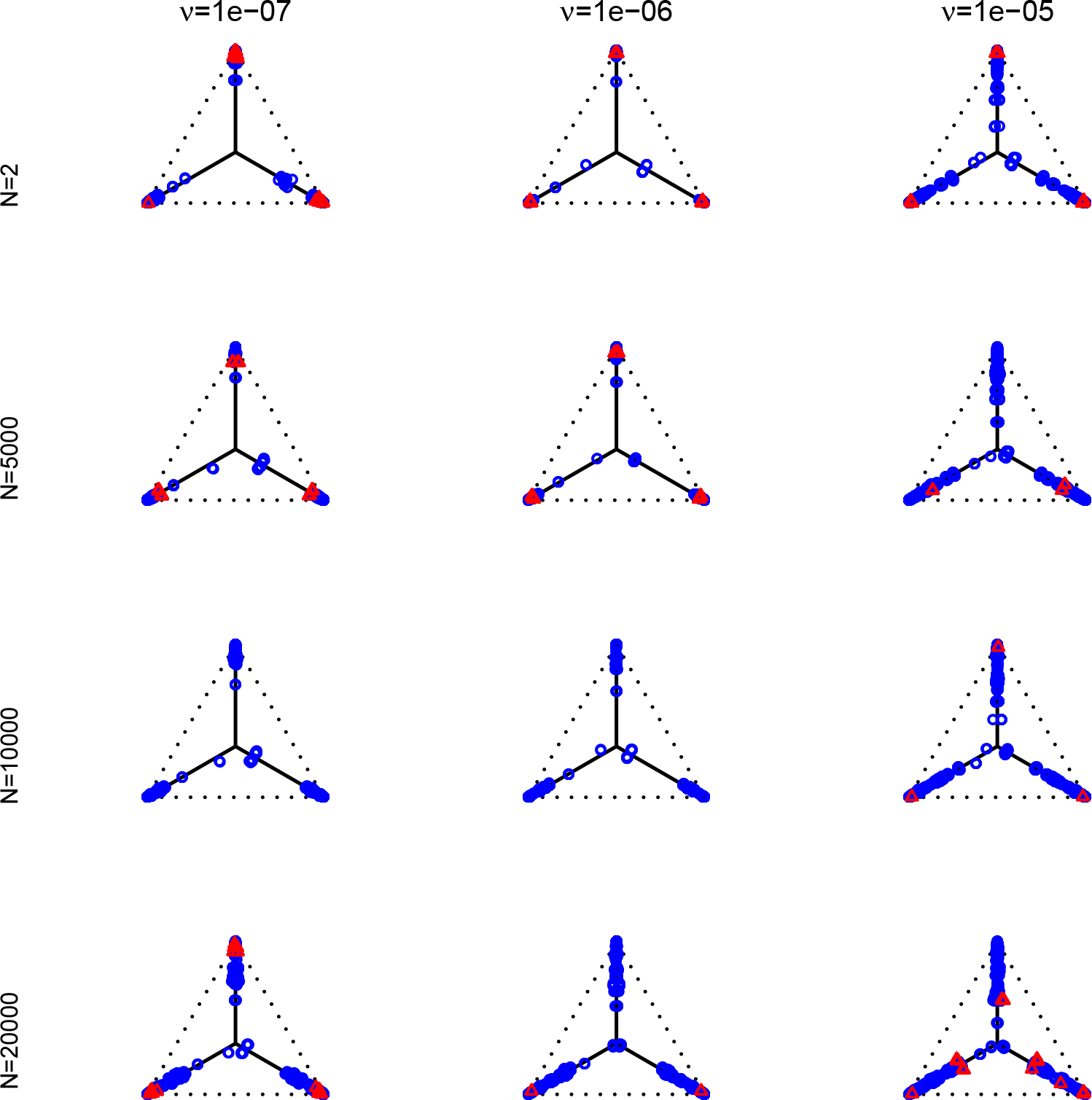
Empirical *CF*s for the MSC on the species tree of Figure 7 are shown for 1000 gene trees inferred from simulated sequences data of length 500 bp. Population sizes for rows are from top to bottom *N* = 2, 5000, 10000, and 20000, and values of *ν* for columns are 10^*a*^ for *a* = −7, −6, −5. Counts of unresolved gene quartets were omitted in this analysis. Red triangles denote empirical *CF*s for which the hypothesis test with level *α* = 0.01 rejects the T3 model, and thus the MSC on a tree.

One important observation from these simulations is that under the ND model, it is possible to simulate data sets and produce a simplex plot that may appear to better match expectations under the MSC model than under the ND. Indeed, in Figure 9, the differences between the upper right plot for *N* = 2*, ν* = 10^−5^ and the lower left plot for *N* = 20000, *ν* = 10^−7^ are not great, even though the first is the ND model with moderate inference error and the second has substantial ILS. A closer look at the data behind these plots, however, reveals more. For the lower left plot the mean raw (Hamming) distance between all pairs of sequences for all genes is ≈ .037 and with this low level of sequence differences, one should expect gene tree inference error to be high. Indeed, virtually all gene trees are inferred with polytomies. For the upper right figure, the mean raw distance is very high, ≈ .664. Here again we expect high gene tree inference error since the sequences have so few sites in agreement. In this case we found about one quarter of the trees had polytomies. While counting polytomies does not fully assess inference error, it is one sign that the quality of a collection of inferred gene trees may be poor.

For *N* = 2 and all parameter choices on this tree that we found led to well-resolved inferred gene trees, the great majority of *CF*s were plotted in the vicinity of the vertices. Nonetheless, we currently lack a clear procedure for distinguishing in these plots between the effects of ILS modeled by the MSC and the ND model with gene tree inference error.

For a final analysis presented in Figure 10, we consider only the set of inferred gene trees for *N* = 10000 and *ν* = 10^−6^, but in counting displayed quartets we treat as unresolved any whose internal branch length was less than *ϵ* for *ϵ* = 10^−*b*^ with *b* = −6, −4, −2. For *b* = −4, −2 this causes more quartets to be treated as unresolved, while for *b* = −6 it has no effect, as the inferred gene trees had a minimum positive branch length of 10^−6^. We then produced simplex plots to investigate the differences between redistributing unresolved quartets as 1/3 of each resolution, and discarding them. The online appendix shows figures of this sort for *N* = 10000 again, but with the more extreme substitution levels from *ν* = 10^−7^ and *ν* = 10^−5^.

**Fig. 10.**
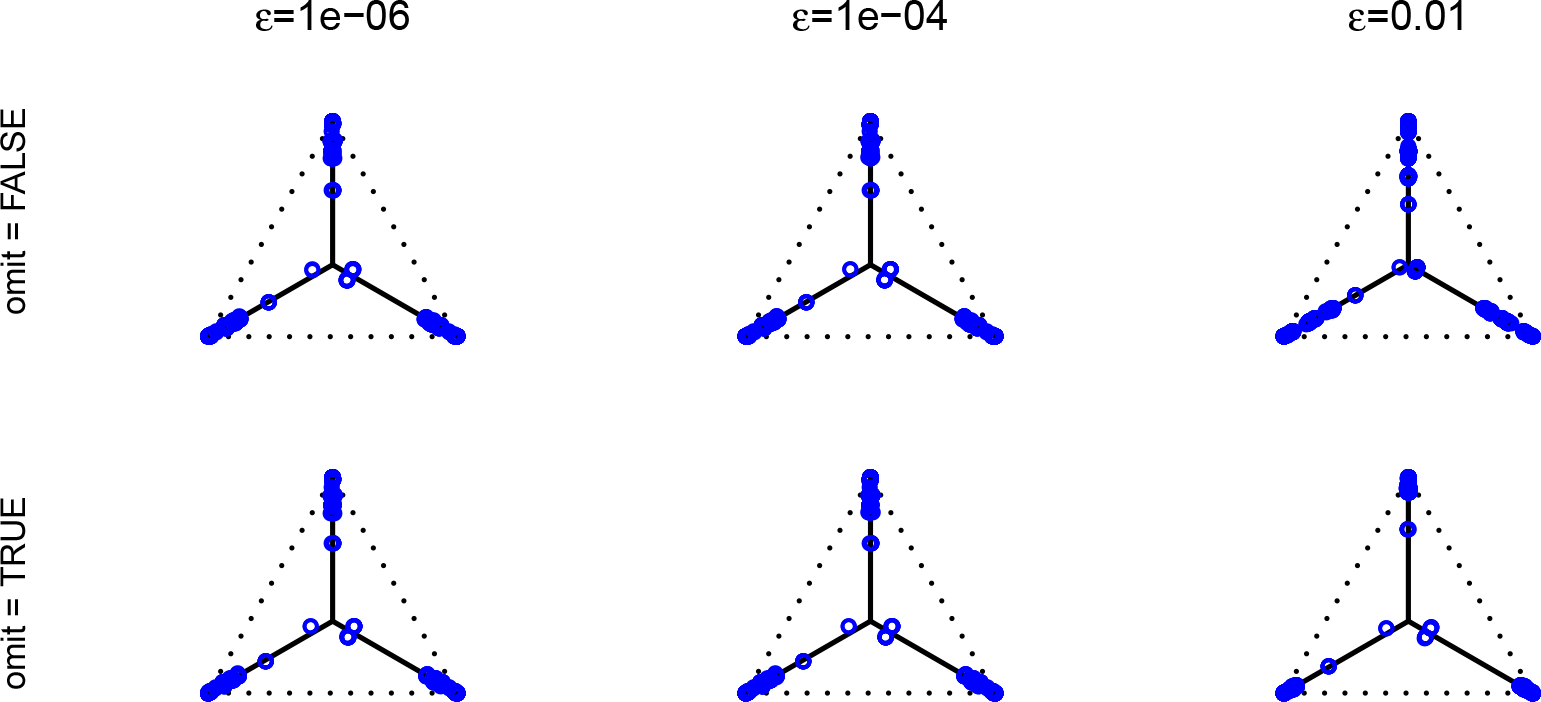
Empirical *CF*s for the MSC on the species tree of Figure 7 are shown for 1000 gene trees inferred from simulated sequences data of length 500 bp, for *N* = 10000 and *ν* = 10^−6^. In columns, displayed quartets with internal edge lengths less than *ϵ* for *ϵ* = 10^−*b*^ for *b* = −6, −4, −2 were counted as unresolved. Counts of unresolved quartets were then either redistributed as 1/3 of each resolution (omit=FALSE) for the top row, or ignored (omit=TRUE) for the bottom row. Lack of red triangles indicates there are no rejections of the null hypothesis for the T3 test with level *α* = 0.01 for these empirical *CF*s.

The choice of *b* = −2 for the right column of Figure 10 is somewhat large for this data set, but is useful for understanding. This value causes so many quartets to be treated as unresolved that the differing effects of the two treatments is noticeable. If such a count is redistributed as 1/3 to each resolution, this tends to equalize the entries of the *CF*, causing substantial movement toward the centroid. On the other hand, if an unresolved count is omitted, much topological discord in the quartets may be removed, and we see the *CF* move toward a vertex.

The two values of *b* = −6, −4 give plots in the first two columns of Figure 10 that are difficult to distinguish visually across either rows or columns. When there are few short edges in the gene trees, this is what one should expect, since the choice of treatment will result in only small changes in counts. However, for low levels of substitutions over the gene trees, this may not be the case, as many gene tree branches may be short. This is illustrated in the appendix in the similar plot for *ν* = 10^−7^. When empirical data is explored through simplex plots we recommend preliminary plots be made with both treatments of unresolved counts, and with a range of length cutoffs for treating quartets as unresolved.

## Results

We illustrate the use of the quartet simplex plots and hypothesis tests described earlier by examining several empirical data sets. The data sets we discuss have appeared in other publications, and we use gene trees previously inferred for those works. Although inference error for these gene trees may have been minimized as much as current methods allow, we expect such error is still present.

### Yeast

The multilocus yeast data set of Rokas et al. [2003] was used in an early attempt to infer a species tree from genomic data. Here we consider a much-analyzed subset of it, with 106 gene trees on 8 taxa, and thus 70 different 4-taxon sets. Both the initial analysis, and many subsequent ones, e.g. [Phillips et al., 2004, Collins et al., 2005, Hedtke et al., 2006, Jayaswal et al., 2014], sought a common tree on which all the loci evolved.

There are some polytomies in the gene trees, and we present an analysis treating unresolved quartets as 1/3 of each resolution. Results are similar if unresolved quartets are discarded. Figure 11a shows a simplex plot and independent T3 hypothesis test results for each *qcCF*, using an initial significance level of 0.05. Momentarily ignoring the color/shape-coded hypothesis test results in the figure, a glance at the simplex plot shows some empirical *CF*s far from the T3 model lines, suggesting the MSC on a species tree is not consistent with these data. However, with only 106 gene trees, one might at first imagine that the deviations we see from the model lines could be due to chance.

**Fig. 11.**
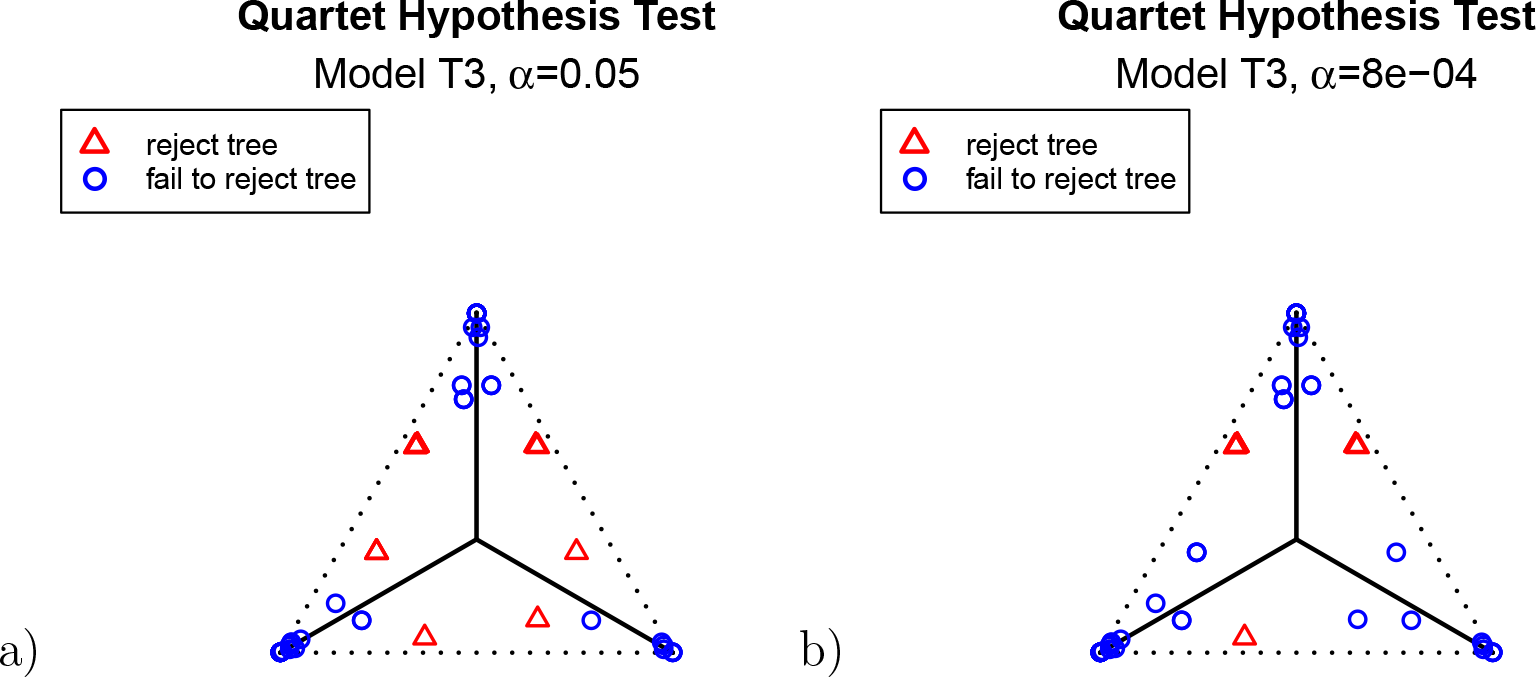
Simplex plots of empirical *CF*s for a Yeast data set of Rokas et al. [2003], composed of 106 gene trees on 8 taxa. While some *CF*s are plotted near the T3 model lines, even without a formal hypothesis test, those far from the T3 model lines suggest a very poor fit of these gene trees to the MSC on a species tree. As described in the text, formal hypothesis tests at the levels *α* = 0.05 and *α* = 8e-4 confirms this is the case for many of the quartets individually. Even when the Holm-Bonferroni method is used to adjust for multiple comparisons, there is strong evidence for rejecting the MSC on a species tree at level *α* = 0.05.

The hypothesis tests, however, allow the number of gene trees to be taken into account in interpreting which *qcCF*s are extreme. With a level of *α* = 0.05, the results of independent tests for each *qcCF* give some red triangles in the plot, further indicating rejection of the MSC on a species tree. However, since these tests are designed to address only sampling error, and gene tree inference error is likely present, a smaller level *α* might be preferred.

A more detailed inspection of *p*-values from the T3 test shows the 9 empirical *CF*s plotted as red triangles closest to the simplex boundary have *p* < 1.11 × 10^−7^, while the remaining 5 plotted as red triangles in Figure 11a have *p* < 8.71 × 10^−4^. The next largest *p*-value, which is associated to a point plotted as a blue circle, is approximately 0.0894. For example, *α* = 8 × 10^−4^ yields the plot of Figure 11b which separates the two most extreme groups. Such plots for several choices of *α* allow viewers to visually distinguish levels of model fit. We strongly caution against over reliance on any single threshold as determining which quartets fit the model and which do not. Data sets of gene trees are likely to be somewhat noisy due to inference error, and this may lead to smaller *p*-values. Nonetheless, the *p*-values provide a standardized way of quantifying ‘extremeness’ across all choices of 4 taxa.

Since the possibility remains that some small *p*-values arose only because we performed so many tests and treated them as independent, we apply the Holm-Bonferroni method, with *α* = 0.05. This leads to rejection of the null hypothesis only for the 9 choices of four taxa giving the smallest *p*-values, whose empirical *CF*s are red triangles in Figure 11b. In fact, even though the Holm-Bonferroni adjustment is quite conservative, a level of *α* ≈ 3 × 10^−6^ or smaller would be needed to avoid all rejections of the null hypotheses that the gene trees arose from the MSC on a species tree.

An alternative viewpoint is that all or most of the gene tree discord in this data set is due to gene tree inference error. There is no principled way to rule out that possibility, in part since model misspecification might have played a role when inferring the gene trees. However, general acceptance of the validity of standard phylogenetic methods argues otherwise. It should be emphasized that the methods introduced here use only unrooted topologies of gene trees, which is considered to be the aspect of gene trees that can be most accurately inferred. At the very least, a claim of such extreme inference error should require a detailed argument as to why gene tree error would be so great for certain quartets and not others. Assuming gene tree inference error is not large, the reasonable conclusion is that this data set is not consistent with the MSC on a species tree, and thus also not consistent with its ND model.

With the MSC on a species tree rejected, one might next consider that the taxa should be related by a network. Indeed, many authors [Bloomquist and Suchard, 2010, Holland et al., 2004, Wen and Nakhleh, 2018, Wu et al., 2008, Yu et al., 2011, Huson et al., 2010] have considered a network as appropriate for these data, and investigated evidence for reticulations. Particularly relevant is the recent inference of a network from these gene trees under the Network MSC model by Allman et al. [2019b] using NANUQ. In that algorithm rejection of the T3 model for a *qcCF* leads to use of a hybridization model for the four taxa, and with all hybrid quartets so identified a network can be quickly inferred.

### Mammals

The multilocus mammal data set of Song et al. [2012] was first analyzed under the MSC using both the rooted triple pseudolikelihood algorithm MPest [Liu et al., 2010] as well as the clade-based distance algorithm STAR [Liu et al., 2009], to obtain a species tree. The conclusions of those analyses were disputed by Gatesy and Springer [2013], who found fault with using the MSC as a basis for species tree inference in the presence of gene tree inference error, preferring a direct analysis of the concatenated sequences. See also the response from Wu et al. [2013].

From the standpoint of model-based inference, this controversy is based on a choice between the ND model of ‘no discord among true gene trees’ and the MSC model of ILS in formation of the gene trees. While hybridization, or any substantial non-tree-like signal in the data set, seems unlikely for mammals, we note that both the ND and MSC models simply ignore the possibility.

Since the original collection of gene trees for the data had several errors, we use those from the re-analysis of Mirarab et al. [2014]. This consists of 424 gene trees on 37 taxa, so there are 66045 choices of four taxa for which the *qcCF*s are tabulated. All of the gene trees are fully resolved. We explored the data set using a range of testing options, including cutoffs as large as 0.01 for internal quartet edge lengths to treat quartets as unresolved, and both discarding or redistributing unresolved counts. Here we present an analysis only for the fully resolved trees, since the other analyses made little difference in conclusions.

The T3 simplex plot on the left in Figure 12a shows that all empirical *CF*s for the data set lie relatively close to the model lines. With a level of *α* = 0.001, which was chosen to accommodate some likely gene tree inference error, the test rejects the MSC on a tree for some quartets. A closer look shows that the smallest *p*-values for any choice of four taxa are around 7.7 × 10^−7^, though this arises for points near the simplex vertices, where the most supported topology is clear. Applying the Holm-Bonferroni method, we find no rejection of tree-likeness even for *α* = 0.05. While this can be a very conservative test, there is no strong evidence for rejecting the MSC on a species tree, a conclusion consistent with other biological knowledge.

**Fig. 12.**
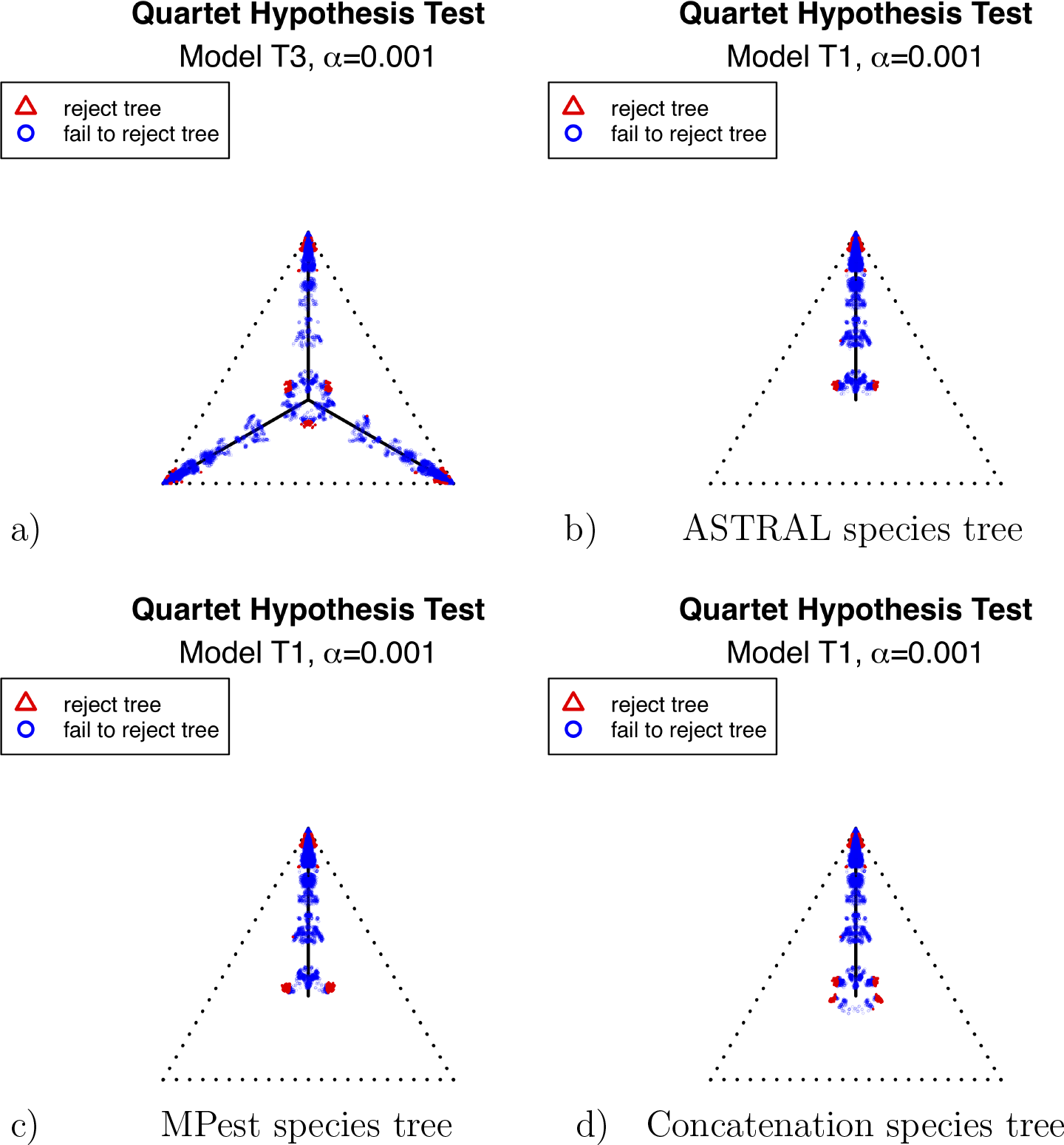
Simplex plots for the Mammal data set of Song et al. [2012], for 424 gene trees on 37 taxa. In (a), model T3 is used, as no species tree is hypothesized. In (b) the putative species tree obtained by ASTRAL, ASTRAL-III, and QDC, is used for determining *CF* placement under the T1 model. In (c), the putative species tree is that inferred by MPest. In (d) the putative species tree is that obtained by a Bayesian concatenation method. All trees are as reported in Mirarab et al. [2014]. The species trees in (b) and (c) differ in the placement of only one taxon, the tree shrew (*Tupaia belanger*i), which is moved by a nearest-neighbor interchange. The tree in (d) differs from that in (c) by an additional NNI move, applied to the Carnivora-Perissodactyla clade. See original publications for more complete descriptions.

We have no formal statistical procedure to decide whether the simplex plot looks as it does because of ILS modeled by the MSC with small or moderate inference error, or because true gene trees arose from the ND model with substantial inference error. Nonetheless, the location of the empirical *CF*s so close to the T3 model lines suggests that the MSC potentially fits the data set well, without a need to assume substantial inference error. Even the clustering of points in the plot is reminiscent of the equality of expected *CF*s for many quartets, as discussed for Figure 4, a feature which tends to lead to similar clusters of empirical *CF*s in simulations. Thus we find ILS a simpler explanation of this plot than the presence of substantial gene tree error mimicking ILS, and suggest a coalescent-based analysis is warranted.

After inferring a species tree topology from the data set by any method one chooses — whether based on the MSC or not — a new simplex plot can be produced hypothesizing the inferred topology for the model T1. For the mammal data we consider three different choices of hypothesized species trees which have appeared in earlier analyses. The trees are estimated by the coalescent-based ASTRAL and MPest, and a non-coalescent-based Bayesian concatenation analysis, as described by Mirarab et al. [2014] (and, in part, for the original data set by Song et al. [2012]). We also inferred species trees by the MSC based methods ASTRAL-III [Zhang et al., 2018] and QDC [Rhodes, 2019], obtaining the same species tree as from ASTRAL.

The MPest tree differs from each of the other two by 1 NNI (nearest neighbor interchange) move, while the ASTRAL and concatenation tree differ from each other by 2 NNIs. The three species trees differ from one another only in the placement of the tree shrew (*Tupaia belangeri*) by an NNI, and of the placement of one clade (composed of Perissodactyla and Carnivora) by an NNI.

In Figure 12b,c,d, simplex plots are given for the T1 model hypothesizing these species trees. The visual differences between Figure 12b (ASTRAL tree) and c (MPest tree), are subtle, involving the placement of the two ‘lobes’ of red triangles located near the centroid. In Figure 12b, these are slightly closer to the model line, suggesting a slightly better fit of the ASTRAL tree than the MPest tree. However, our plots and tests are not designed to measure fit to a single species tree, but rather the fit of each *qcCF* independently to the induced quartet tree.

The simplex plot in Figure 12d for the concatenation tree shows more interesting differences from those of 12b,c. Although the concatenation tree differs from the MPest tree by only an NNI move applied to a single clade, this affects where a relatively large number of empirical *CF*s are plotted, in a ‘ring’ around the centroid. While gene trees sampled directly from the MSC on a species tree with a near 0 length branch can lead to a cloud of *CF*s around the centroid, a ring does not arise. Thus plausible conjectures are either this species tree is not correct, or that the gene tree inference procedure led to a bias away from equal occurrences of the 3 quartet topologies when the true quartet gene trees had nearly equal frequencies.

Investigating individual *p*-values for the T1 tests for the three species trees gives additional insight, but no firm conclusions. For all three, the smallest *p*-values arose for quartets whose empirical *CF* is plotted near the top vertex of the simplex, where the most supported topology is clear. The proportion of quartets for which the hypothesized species tree topology was rejected is shown in Table 1, with these numbers perhaps suggesting a slightly better fit for the ASTRAL tree. Between the two different values of *α* in that table, though, whether the MPest or concatenation tree is rejected more frequently switches. (Note that theory does not suggest that the rejection fraction should be close to *α*, since the tests are not of independent hypotheses. Moreover, since we expect some noise in the inferred gene trees, the fraction rejected is likely to be inflated over what we would see with true gene trees.)

**Table 1.** For the mammal gene trees, the proportion of *qcCF*s for which the model T1 determined by a hypothesized species tree leads to rejection at the level *α*, rounded to 4 decimal places.

| species tree | $\alpha = .05$ | $\alpha = .001$ |
| --- | --- | --- |
| ASTRAL | 0.1108 | 0.0088 |
| MPest | 0.1109 | 0.0189 |
| Concatenation | 0.1191 | 0.0132 |

One might still argue that all discord in these gene trees is due to inference error, and adopt concatenation as the only justified analysis. We believe that a combination of the MSC and only moderate inference error provides a more parsimonious explanation of the data. However, from the perspective of the quartet simplex plots and hypothesis tests, there is no definitive evidence in favor of one of the three trees over the others. While one might rank them in the order of b,c,d from ‘best’ to ‘worst’ fit, the differences these methods reveal are quite minor. Indeed, the conservative Holm-Bonferroni method confirms this by failing to reject the MSC on any of these trees at the level *α* = 0.05.

### Other examples

We briefly show results from applying simplex plots and hypothesis testing to a few other multilocus data sets. Solís-Lemus and Ané [2016] applied their SNAQ software to a data set of 1183 fully-resolved gene trees inferred from sequences collected from 24 fish species first presented by Cui et al. [2013], to infer network relationships. The choice of a network analysis was based on a high level of discord among the gene trees. A simplex plot for the quartets displayed on the gene trees immediately communicates this, as shown in Figure 13a. The smallest *p*-values for the T3 tests are around 4.7 × 10^−69^, and the Holm-Bonferroni method with *α* = 0.05 leads to rejection of the null hypothesis for 1815 of 10626 tests, about 17%. If the null hypothesis is true, we expect at least one false rejection at most *α* = 0.05 of the time. Since the MSC on a tree cannot explain the data, one must conclude there is either a huge amount of gene tree inference error, or some more complicated model such as hybridization is needed for analysis.

**Fig. 13.**
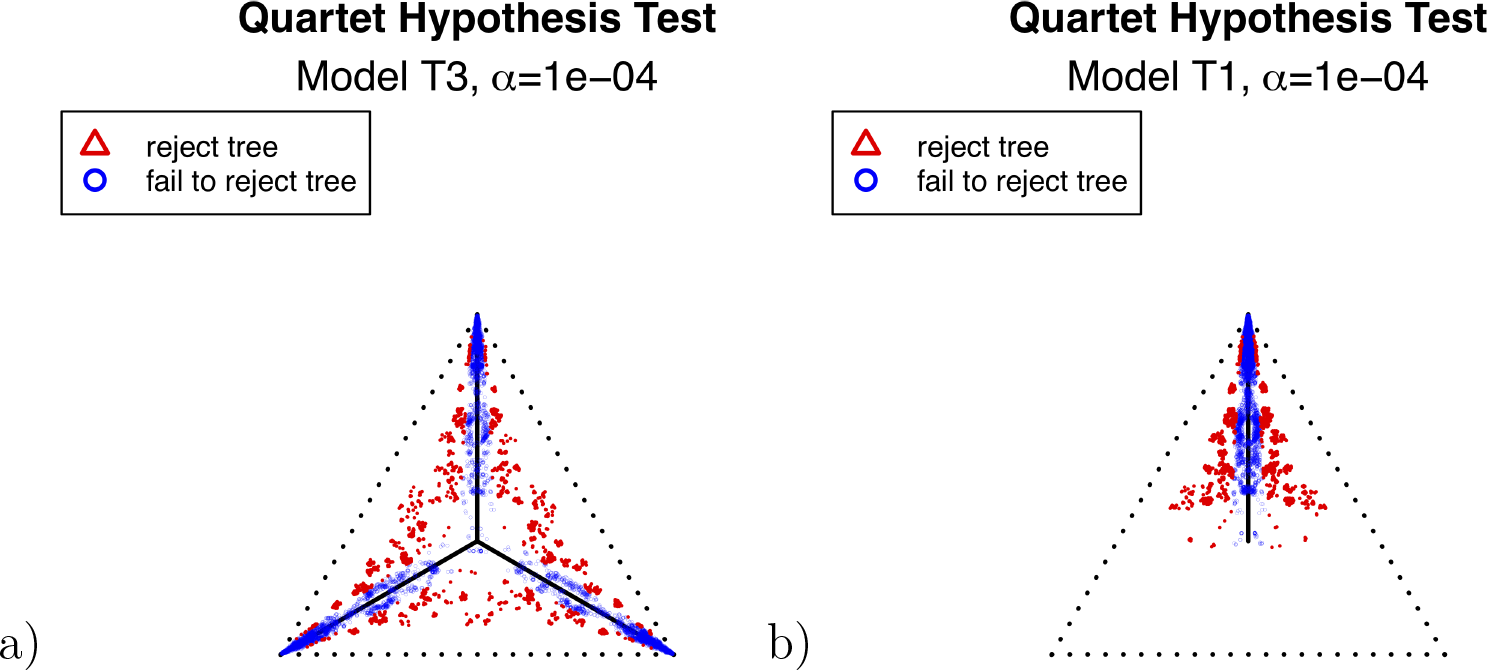
Simplex plots for the Fish data of Cui et al. [2013], composed of 1183 fully-resolved gene trees on 24 taxa. In (a), no species tree is hypothesized. In (b) the putative species tree inferred by ASTRAL-III and QDC, is used for determining *CF* placement under the T1 model.

In spite of this finding, we applied ASTRAL-III and QDC to the gene trees, obtaining the same inferred tree by both methods. Figure 13b shows a simplex plot for the model T1 with this hypothesized species tree. The plotted points and *p*-values still suggest a need for analysis with a more complicated model, such as one capable of inferring a network, but interestingly we see that almost all plotted points are closer to the T1 model line segment than to line segments from other vertices to the centroid. This suggests that there is a ‘dominant’ tree like signal, as *CF*s arising from hybridization on level-1 networks are convex combinations of those on the trees displayed on the network [Solís-Lemus and Ané, 2016, Baños, 2019], and placement of points in the T1 simplex plot above shows that the inferred tree here would be displayed in the full network.

Martins et al. [2014] investigated a multilocus *Drosophila* data set of 4591 gene trees, relating 12 subspecies but with taxa missing from some gene trees. These trees include only single-copy gene families (orthologs), though the full data set included additional gene trees with paralogs. These trees were used in the original publication to benchmark a method of tree inference under a model of gene tree discordance allowing for both ILS and gene duplication and loss. In particular, this is a data set believed to have evolved by processes not modeled by the MSC on a tree.

In Figure 14a and b we show T3 and T1 simplex plots for these gene trees, where the hypothesized (sub-)species tree is that inferred by ASTRAL-III and QDC. While polytomies were present among the gene trees, and this plot was made by redistributing unresolved quartet counts, there are no qualitative changes if those counts are instead discarded. With the tight clustering of points along the model lines, at first one might imagine the data is well described by the MSC, or even (since plotted points are relatively close to the vertices) result from the ND model with gene tree inference error. Indeed, this is a possibility, since unmodeled gene tree inference error may confound all our methods. However, the smallest *p*-values for the T3 test are around 4 × 10^−13^, and the Holm-Bonferroni method rejects the null hypothesis for some tests even with *α* = 2.1 × 10^−10^. When *α* = 0.05 it rejects the null hypothesis for 74 of 495 tests, about 15% of the tests. Thus a case can be made for processes other than ILS here, assuming gene tree error is not substantial.

**Fig. 14.**
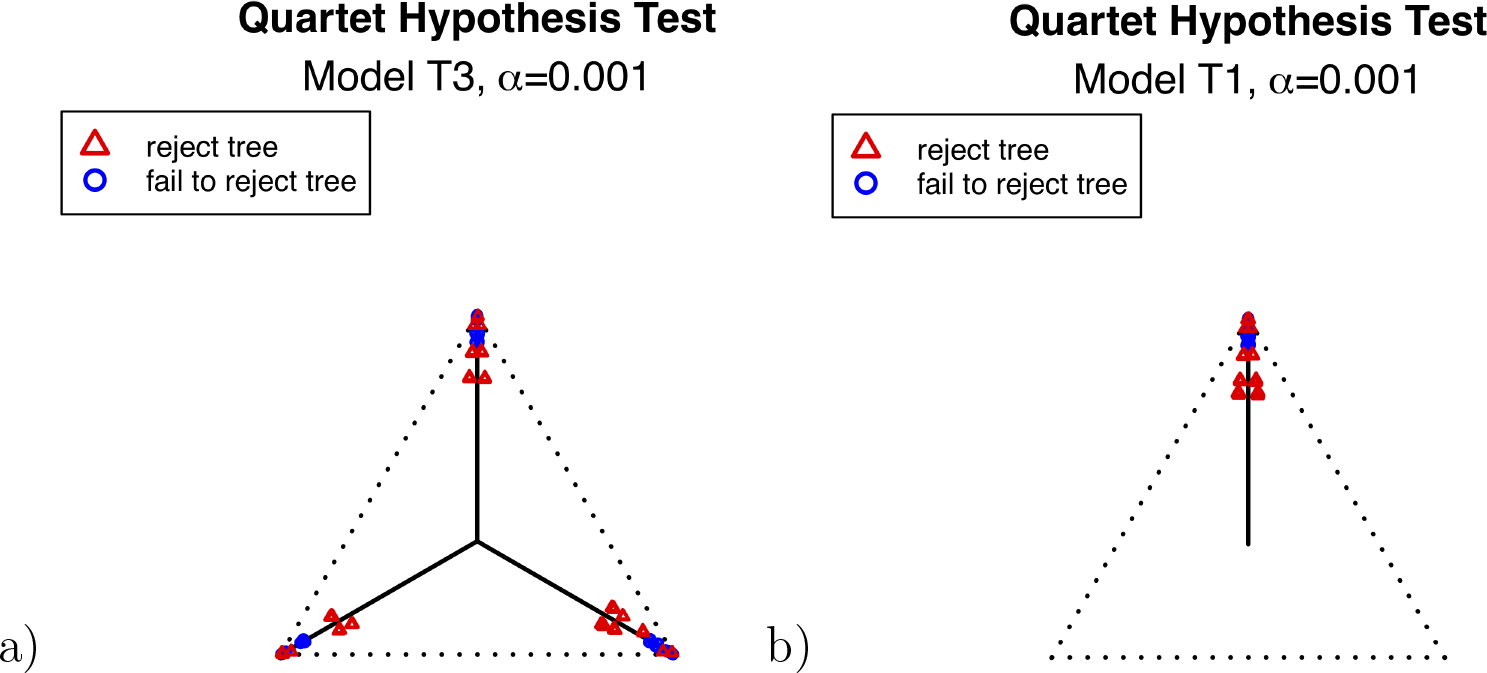
Simplex plots for the *Drosophila* data of Martins et al. [2014], composed of 4591 gene trees on 12 taxa, with unresolved quartet counts redistributed as resolved. In (a), no species tree is hypothesized. In (b) the putative species tree obtained by ASTRAL-III and QDC, is used for determining *CF* placement under the T1 model.

## Discussion

Simplex plots of quartet concordance factors offer a concise and easy-to-understand summary of discordance across gene trees. When such plots show the *CF*s clustered in the corners of the simplex or along the T3 model lines, a researcher may gain insight as to whether the data set might be most productively analyzed under the ‘no discord’ model or the MSC model of ILS. When the points fail to cluster as expected under these models, that indicates either substantial gene tree inference error, or the need for an analysis with a more complicated model, such as one capturing hybridization, or gene duplication and loss.

After a species tree is inferred from either gene trees or the original sequence data, by any method the researcher chooses, a new simplex plot under the T1 model hypothesizing the inferred species tree similarly illustrates gene tree discord with that species tree. Under the MSC, plotted *CF*s should cluster along the T1 model line, while under the ‘no discord’ model (with gene tree inference error) they should be near the top vertex.

The formal hypothesis tests outlined here can indicate how closely a quartet count concordance factor *qcCF* fits the T1 or T3 model, and rejection of these models implies rejection of the MSC, and its ND submodel, on a species tree. Although these tests fail to account for inference error in gene trees, simulations show that moderate gene tree inference error generally manifests itself as a reduction in the *p*-values, but not in extreme changes in the location of plotted concordance factors away from the model lines. In more detail, when gene trees sampled under the MSC are replaced by inferred trees from sequences simulated on them, we see only a moderate spreading of plotted *CF*s around the T1 and T3 model lines. While this phenomenon is visually quite clear, it does make precise interpretation of *p*-values for empirical data sets difficult.

Because of the ubiquitous presence of gene tree inference error, with different gene tree sets possibly having different levels of such error, we caution against using a rigid choice of a level *α* as a meaningful cutoff for rejection for the hypothesis tests. Nonetheless, *p*-values are still valuable. They offer a measure of model fit for a *qcCF* that can be interpreted relative to others from the same collection of gene trees, all of which can be presumed to reflect similar levels of error. In this way we can draw meaningful distinctions between those *qcCF*s that may reflect tree-like species relationships under the MSC, and those suggesting more complex processes. Moreover, as was illustrated by the mammal data set, for some empirical data sets whatever gene tree inference error is present may have minimal impact on the tests outlined here. With either very high or very low levels of sequence dissimilarity, however, one must be concerned about the quality of the gene tree inference, as even the choice of inference software may affect results.

The work of Stenz et al. [2015] is in many aspects close in spirit to what has been presented here, in that it develops tools for measuring fit to a species tree (fully resolved or not) of quartet concordance factors. It differs, however, in the important aspect that it requires a hypothesized metric species tree, while the tests here can be performed without specifying any particular topology (the T3 test), or by specifying a tree topology without branch lengths (the T1 test). In particular this can be especially useful early in an analysis, when it is unclear whether a tree model is even reasonable for the data. Liu et al. [2019] also suggested a hypothesis test, but only for judging which of 2 metric species trees better fits a gene tree collection, and not for whether the MSC on a species tree should itself be rejected.

An interesting feature of the methods of Stenz et al. [2015] is its use of Dirichlet distributions to model the spread of empirical concordance factors around their ‘true’ values. This can be viewed as a way of modeling unknown noise, so that their estimation of the Dirichlet concentration parameter for a data set might provide some measure of combined sampling and inference error. Whether this approach, or a similar one, can be developed to more explicitly address gene tree error in our framework remains a project for the future.

The hypothesis tests presented here were designed to test the MSC model for a single quartet count concordance factor. While the Holm-Bonferroni method provides one way to apply the test to data for larger trees, it is a conservative approach, giving low power to reject the quartet tree models. This may be acceptable if a researcher is comfortable setting a high bar for suspecting non-tree-like species relationships. However development of better methods for trees with more than four taxa is certainly needed, though not straightforward. For instance, if several sets of four taxa have the same internal edge in the quartet tree displayed on the species tree, then their expected *CF*s are identical, and even with a partial overlap of internal quartet edges they are related. The current tests ignore this fact. Similarly, tests for polytomies on *n*-taxon trees and individual edges under the MSC model are also needed.

With implementations of simplex plotting and hypothesis testing tools in the `MSCquartets` R package, the methods described here are available for use in both exploring multilocus data sets, and presenting final analyses. In particular a single simplex plot can communicate much about the level of discord in a collection of gene trees, or of their discord with a hypothesized species tree, allowing readers to form their own judgements of the fit of a data set to an inferred tree. We agree with the statement by Stenz et al. [2015] that it “should become standard practice to conduct tests of the population tree model as part of … an analysis.” Although there is still much room for improving methodology for such tests, simplex plots of quartet concordance factors provide a very useful beginning.

## Acknowledgements

Hector Baños contributed to many fruitful discussions that aided in developing both the methods presented here, and their interpretation. We also thank Jeremy Brown, Rob Lanfear, and Bob Thomson for their organization of the Symposium at which this work was initially presented.

All authors contributed equally to this work.

## APPENDIX Simulation details

The 15-taxon species tree given in Figure 7 of the body of the paper was generated by a pure birth process by the R package ape [Paradis and Schliep, 2018], and then rescaled by a constant so that branch lengths should be interpreted as number of generations. In Newick notation, the species tree is *σ* =

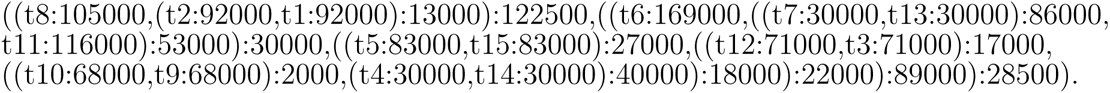

For the substitution process for sequence simulation along sampled gene trees, we chose a GTR model with parameters previously used in several other simulation studies [Chifman and Kubatko, 2014].

In choosing values of *ν*, the conversion factor from units of generations on gene trees to substitution units, we first tried a range of values in preliminary simulations, observing that at both extremes of the range when 500bp sequences were simulated on the sampled gene trees with parameter choices described below, inferred trees tended to be very poorly resolved, due to either excessive similarity or dissimilarity of the sequences. We constructed a plot of the range between maximum and minimum raw (Hamming) distance between simulated sequences on a gene tree, as shown in Figure 15, to choose the range of *ν* values between 10^−7^ and 10^−5^ we used. This 100-fold range in mutation rates results in raw distances encompassing those we computed from several empirical data sets of multilocus sequences for which gene trees have been inferred by other researchers.

**Fig. 15.**
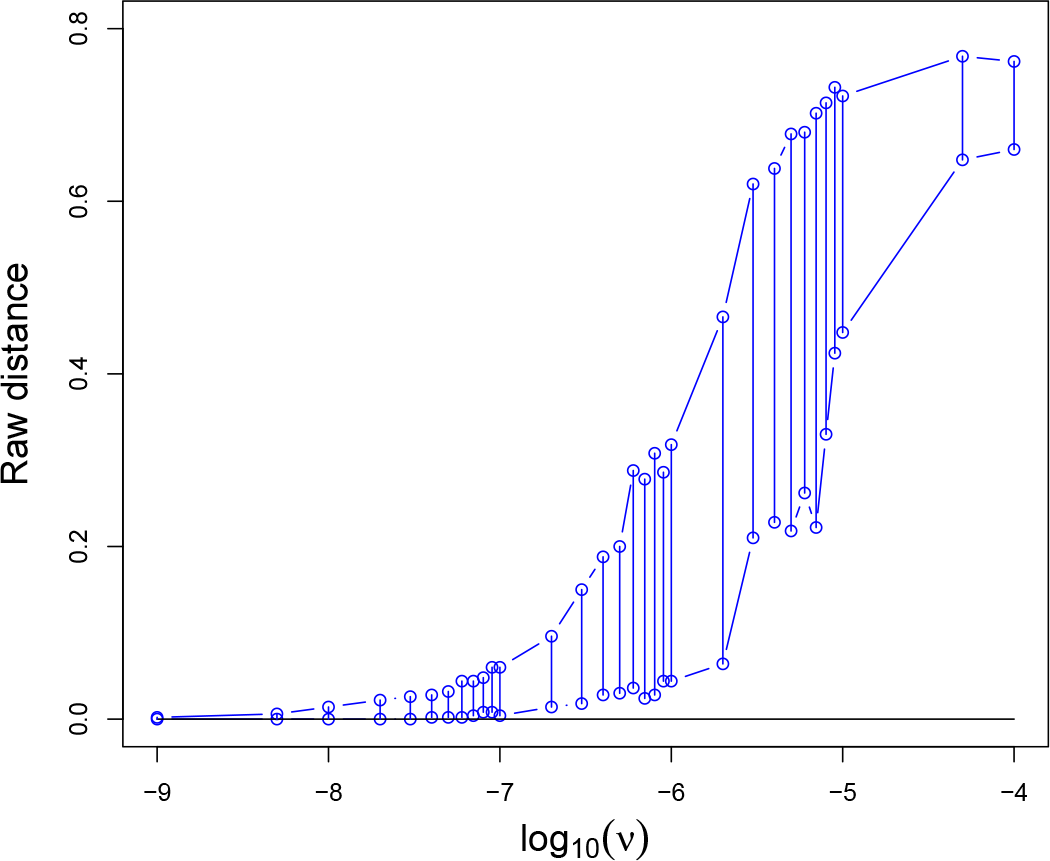
Range of raw (Hamming) distance between sequences simulated on a gene tree sampled from the MSC on *σ* with *N* = 10000 for various values of *ν*. If *ν* < 10^−7^ sequences tend to be very close, while if *ν* > 10^−5^ they tend to be far apart. Either extreme results in poorly resolved or unreliable inferred gene trees.

When inferring gene trees from simulated sequences, we used IQ-TREE with the GTR model, and the `czb` option to contract any inferred short edges. Given our sequence length, this had the effect of contracting edges of length < 10^−6^, to produce polytomies. Our simulation steps were then

1. Using `SimPhy`, for each *N* = 2, 5000, 10000, 20000 and each *ν* = 10^−*a*^ for *a* = −7, −6, −5 simulate 1000 gene trees under the MSC on the tree *σ*. This gives 12 collections of sampled gene trees.
2. For each of the collections of sampled gene trees,

a. For each sampled gene tree, using `SeqGen` simulate aligned sequences of length 500bp on the 15 taxa using the above choices of GTR parameters.
b. For each simulated set of aligned sequences, use `IQ-TREE` to infer a gene tree, using the `czb` option. This is a sampled gene tree with inference error.

This gives 12 collections of sampled gene trees with inference error.

## Additional Simulation Results

Analogs of Figure 9 of the main body of this article are given in Figures 16 and 17, with the number of gene trees reduced from 1000 to 100 and 500 respectively. For these, counts of unresolved quartets are discarded. Figures 18, 19, and 20 show similar plots for 100, 500, and 1000 gene trees with unresolved quartets treated as 1/3 of each resolution.

Figures 21 and 22 show analogs of Figure 10 of the main body of the paper, but for the more extreme low and high levels of substitutions resulting from *ν* = 10^−7^ and *ν* = 10^−5^.

**Fig. 16.**
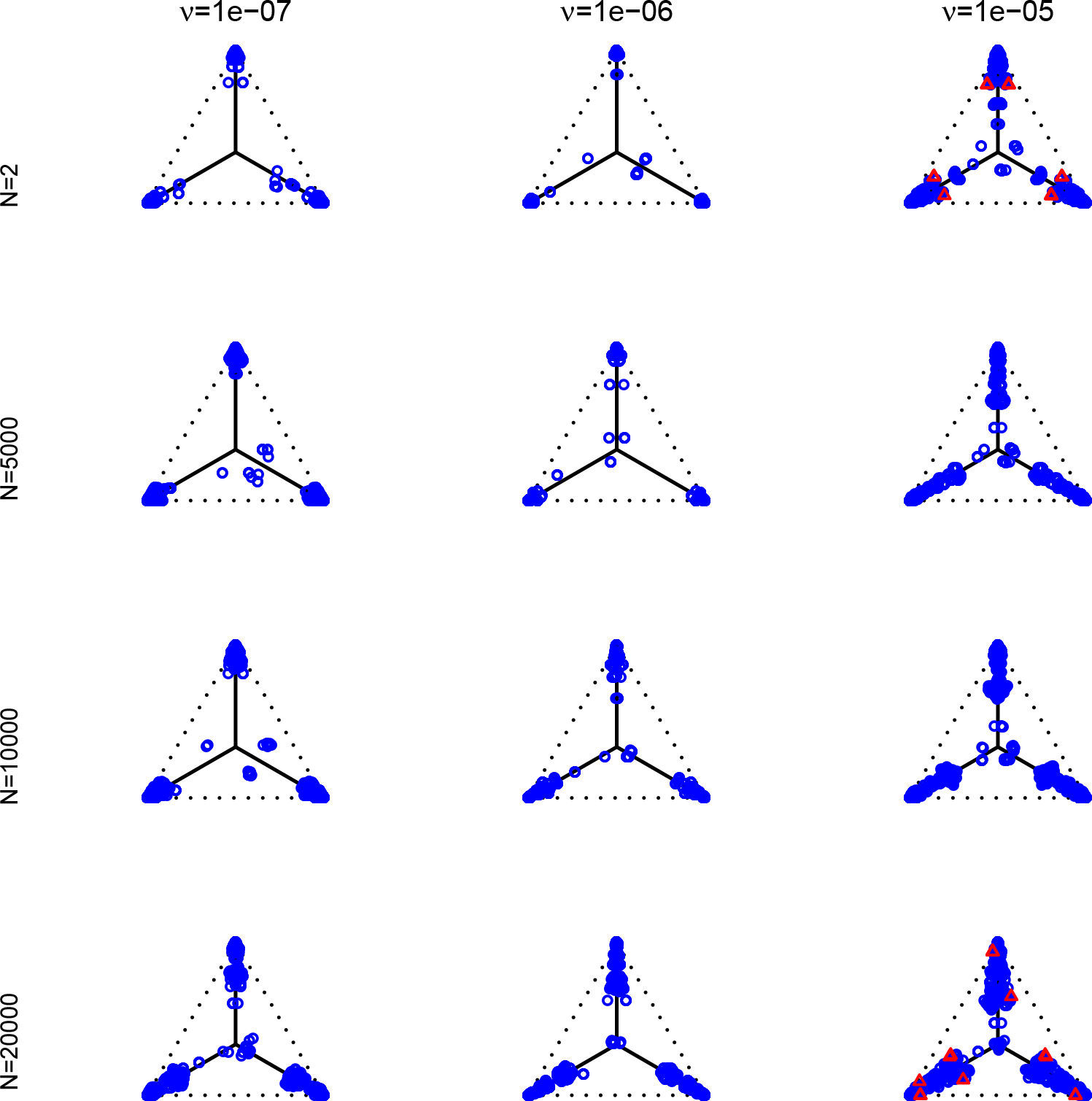
Empirical *CF*s for the MSC on the species tree of Figure 7 are shown for 100 gene trees inferred from simulated sequences data of length 500 bp. Population sizes for rows are *N* = 2, 5000, 10000, and 20000, and values of *ν* for columns are 10^*a*^ for *a* = −7, −6, −5. Counts of unresolved gene quartets were omitted in this analysis. Red triangles denote empirical *CF*s for which the hypothesis test with level *α* = 0.01 rejects the T3 model, and thus the MSC on a tree.

**Fig. 17.**
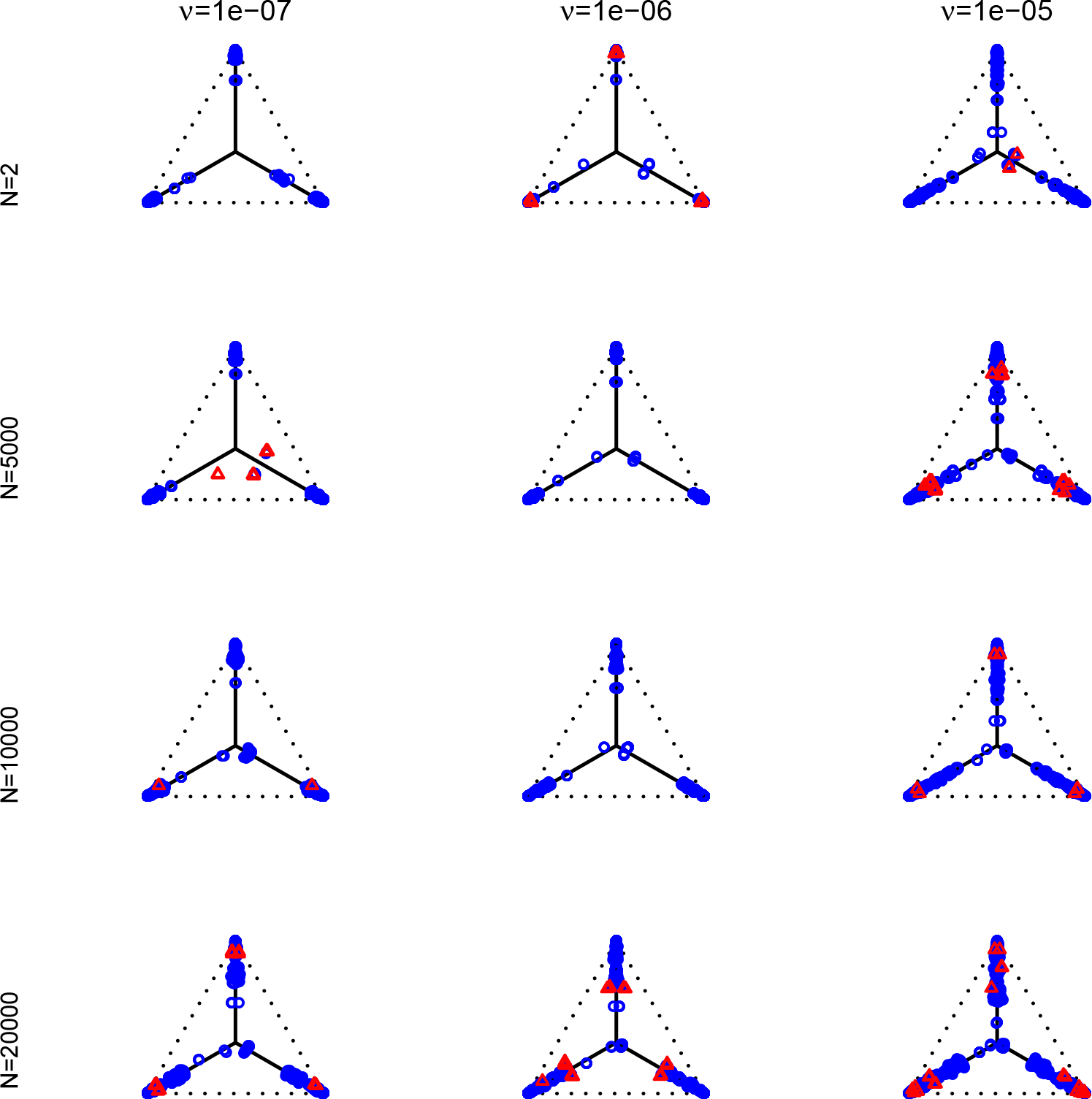
Empirical *CF*s for the MSC on the species tree of Figure 7 are shown for 500 gene trees inferred from simulated sequences data of length 500 bp. Population sizes for rows are *N* = 2, 5000, 10000, and 20000, and values of *ν* for columns are 10^*a*^ for *a* = −7, −6, −5. Counts of unresolved gene quartets were omitted in this analysis. Red triangles denote empirical *CF*s for which the hypothesis test with level *α* = 0.01 rejects the T3 model, and thus the MSC on a tree.

**Fig. 18.**
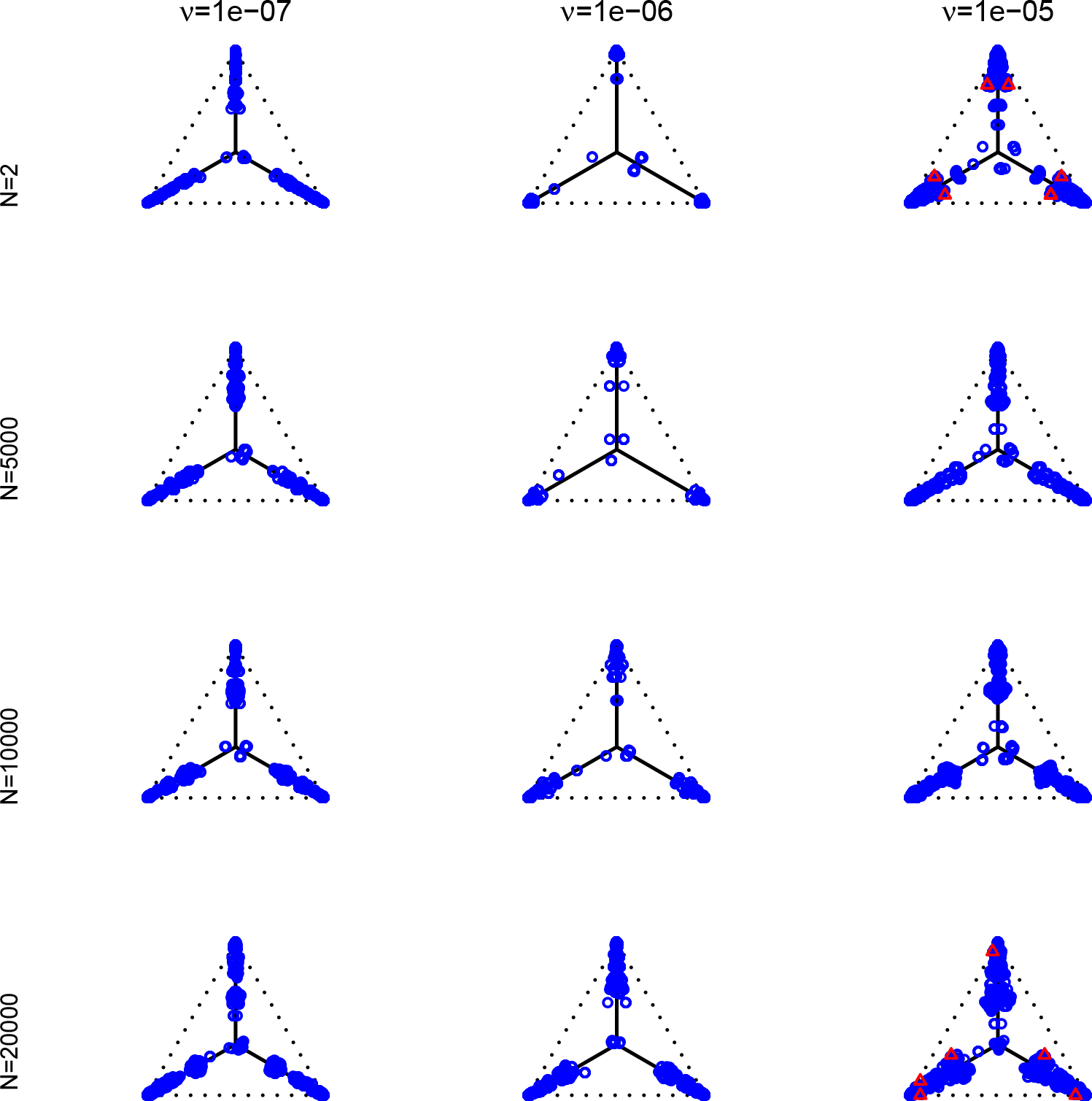
Empirical *CF*s for the MSC on the species tree of Figure 7 are shown for 100 gene trees inferred from simulated sequences data of length 500 bp. Population sizes for rows are *N* = 2, 5000, 10000, and 20000, and values of *ν* for columns are 10^*a*^ for *a* = −7, −6, −5. Each unresolved gene quartet was treated as 1/3 of each resolution. Red triangles denote empirical *CF*s for which the hypothesis test with level *α* = 0.01 rejects the T3 model, and thus the MSC on a tree.

**Fig. 19.**
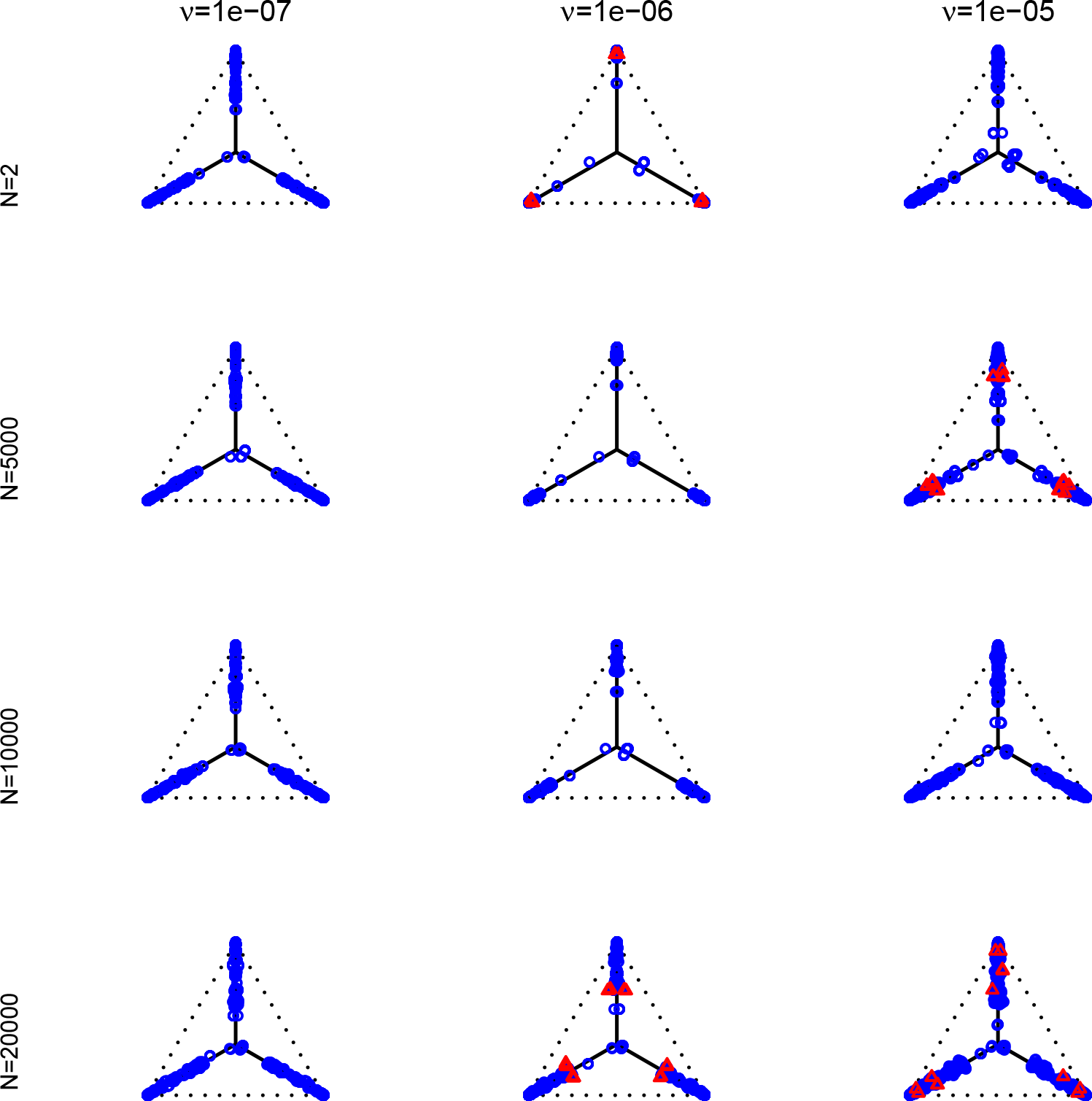
Empirical *CF*s for the MSC on the species tree of Figure 7 are shown for 500 gene trees inferred from simulated sequences data of length 500 bp. Population sizes for rows are *N* = 2, 5000, 10000, and 20000, and values of *ν* for columns are 10^*a*^ for *a* = −7, −6, −5. Each unresolved gene quartet was treated as 1/3 of each resolution. Red triangles denote empirical *CF*s for which the hypothesis test with level *α* = 0.01 rejects the T3 model, and thus the MSC on a tree.

**Fig. 20.**
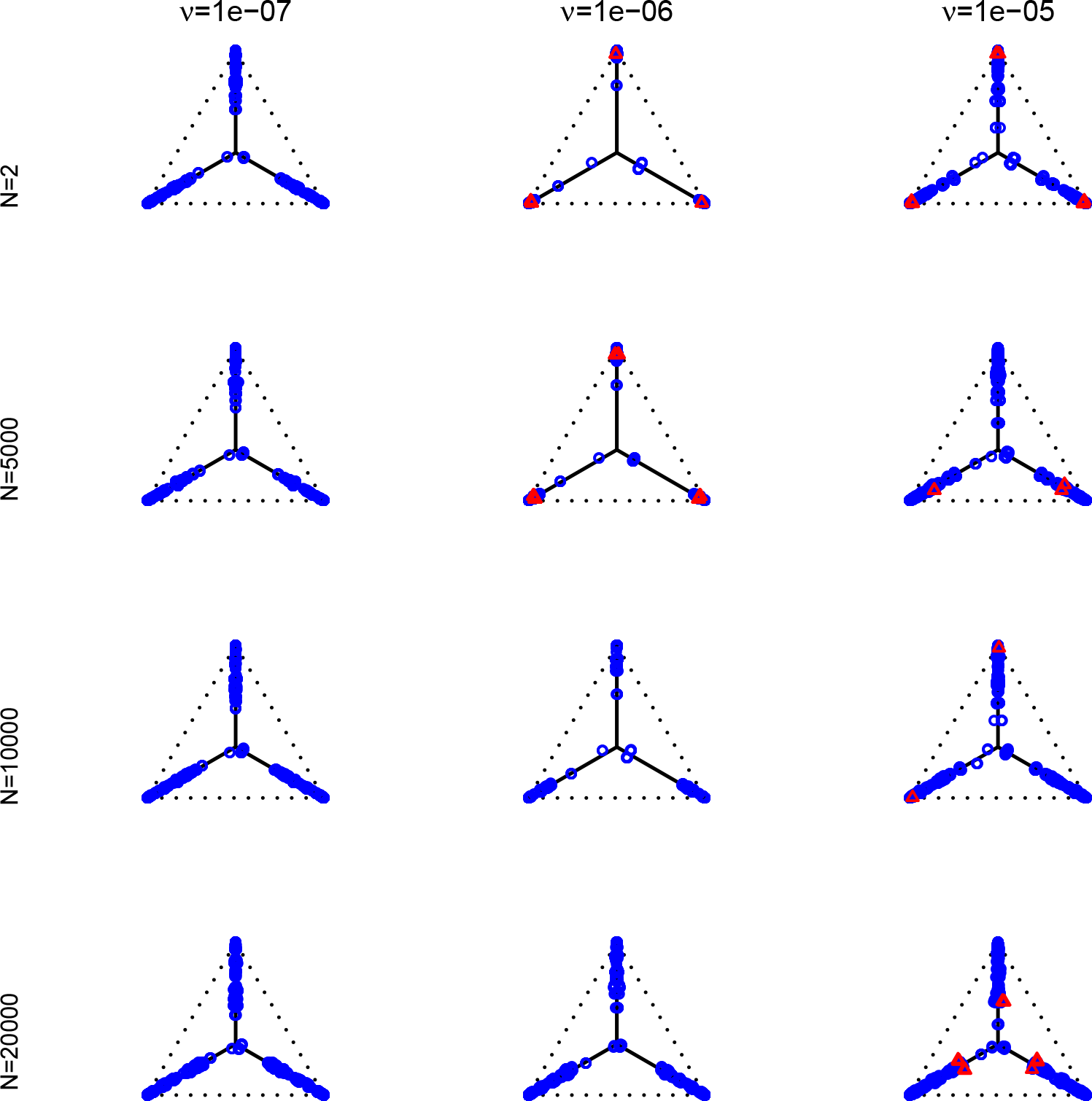
Empirical *CF*s for the MSC on the species tree of Figure 7 are shown for 1000 gene trees inferred from simulated sequences data of length 500 bp. Population sizes for rows are *N* = 2, 5000, 10000, and 20000, and values of *ν* for columns are 10^*a*^ for *a* = −7, −6, −5. Each unresolved gene quartet was treated as 1/3 of each resolution. Red triangles denote empirical *CF*s for which the hypothesis test with level *α* = 0.01 rejects the T3 model, and thus the MSC on a tree.

**Fig. 21.**
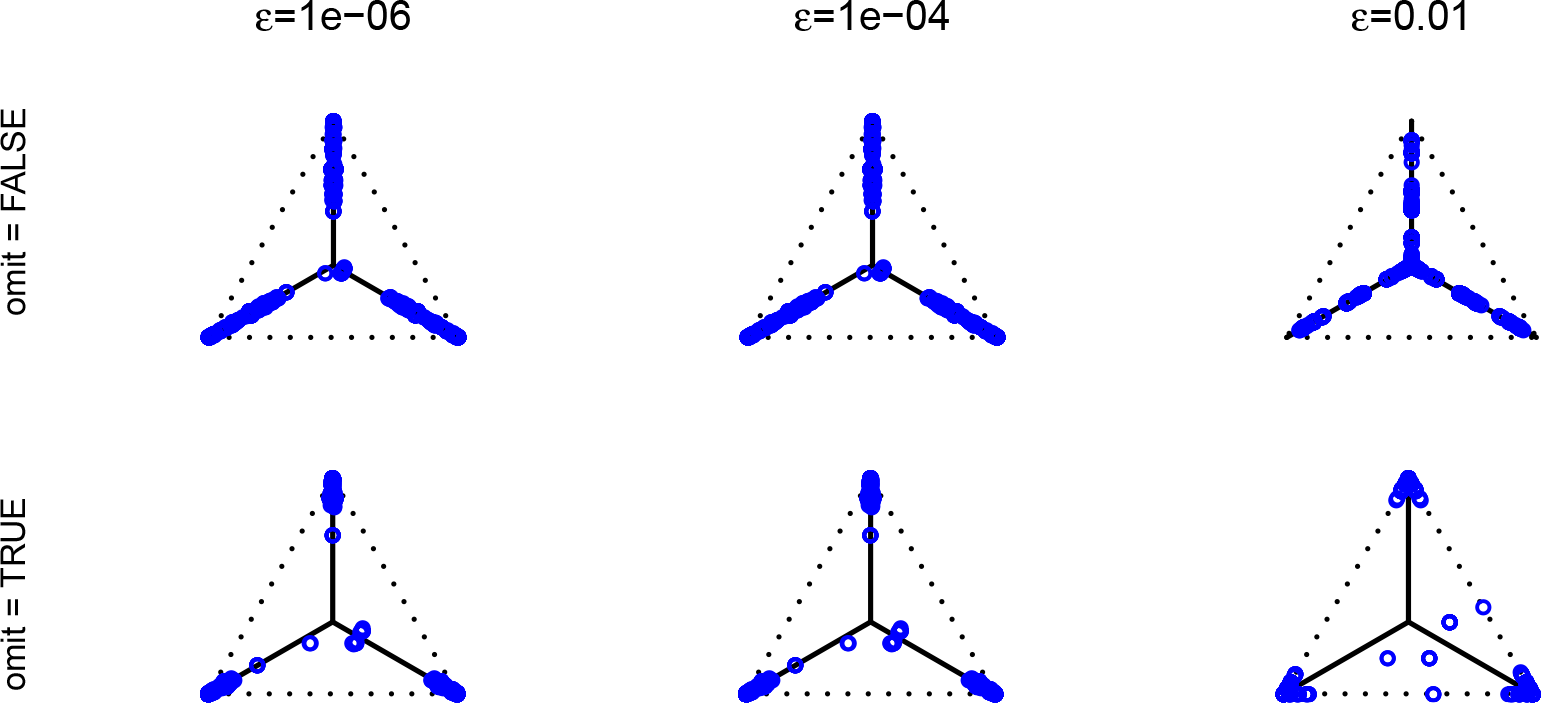
Empirical *CF*s for the MSC on the species tree of Figure 7 are shown for 1000 gene trees inferred from simulated sequences data of length 500 bp, for *N* = 10000 and *ν* = 10^−7^. In columns, displayed quartets with internal edge lengths less than *ϵ* for *ϵ* = 10^−*b*^ for *b* = −6, −4, −2 were counted as unresolved. Counts of unresolved quartets were then either redistributed as 1/3 of each resolution (omit=FALSE) for the top row, or ignored (omit=TRUE) for the bottom row. Lack of red triangles indicates there are no rejections of the null hypothesis for the T3 test with level *α* = 0.01 for these empirical *CF*s. Due to the low total level of substitutions for this data, inferred gene trees have many short branches, so the difference between plots in the top and bottom rows is pronounced. *CF*s far from the model lines in the lower right figure have such low total counts due to omission of unresolved quartet counts that the null hypothesis is not rejected.

**Fig. 22.**
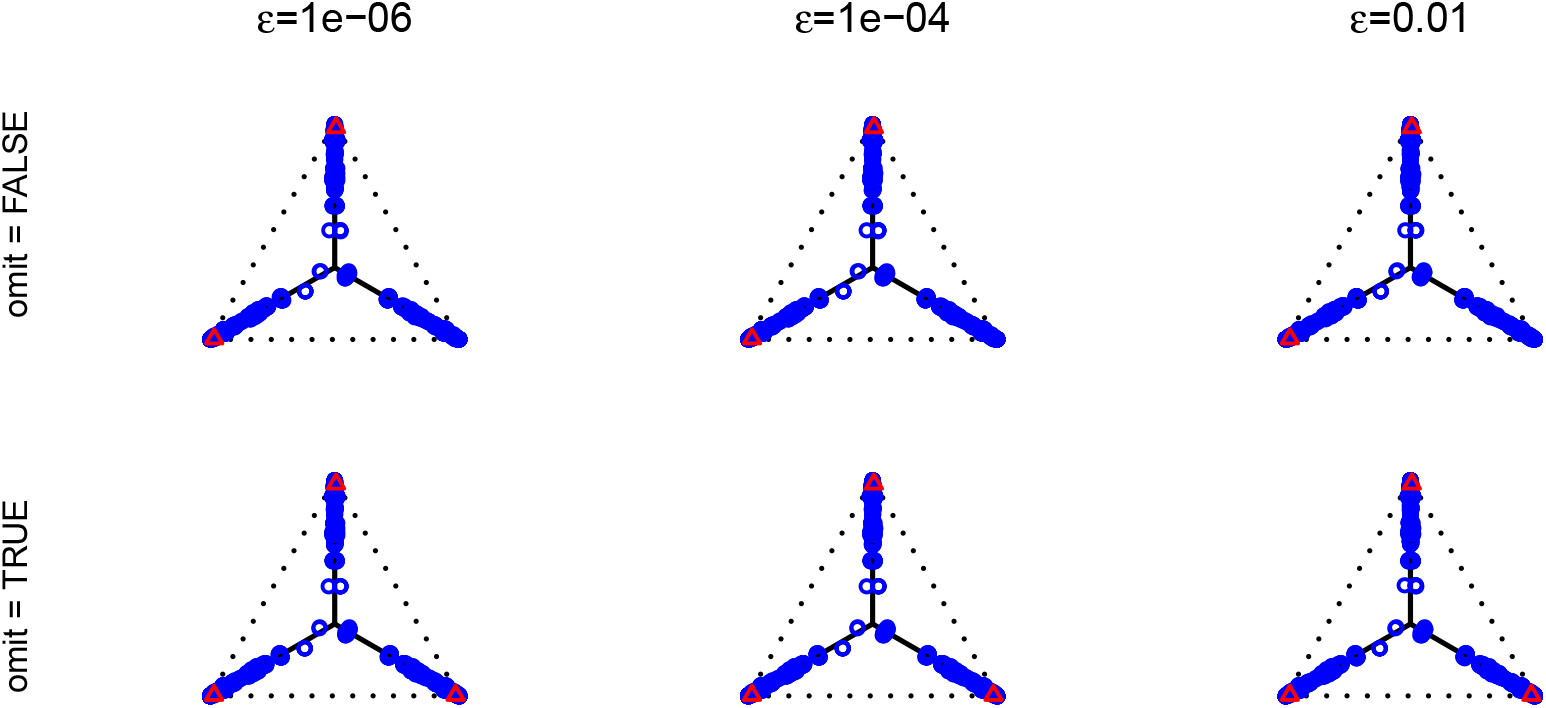
Empirical *CF*s for the MSC on the species tree of Figure 7 are shown for 1000 gene trees inferred from simulated sequences data of length 500 bp, for *N* = 10000 and *ν* = 10^−5^. In columns, displayed quartets with internal edge lengths less than *ϵ* for *ϵ* = 10^−*b*^ for *b* = −6, −4, −2 were counted as unresolved. Counts of unresolved quartets were then either redistributed as 1/3 of each resolution (omit=FALSE) for the top row, or ignored (omit=TRUE) for the bottom row. Red triangles indicate empirical *CF*s for which the hypothesis test with level *α* = 0.01 rejects the T3 model, and thus the MSC on a tree. Due to the high total level of substitutions for this data, inferred gene trees have few short branches, so there is little difference between plots in the top and bottom rows.

